# Efficient Manipulation and Generation of Kirchhoff Polynomials for the Analysis of Non-equilibrium Biochemical Reaction Networks

**DOI:** 10.1101/868323

**Authors:** Pencho Yordanov, Jörg Stelling

## Abstract

Kirchhoff polynomials are central for deriving symbolic steady-state expressions of models whose dynamics are governed by linear diffusion on graphs. In biology, such models have been unified under a common linear framework subsuming studies across areas such as enzyme kinetics, G-protein coupled receptors, ion channels, and gene regulation. Due to “history dependence” away from thermodynamic equilibrium these models suffer from a (super) exponential growth in the size of their symbolic steady-state expressions and respectively Kirchhoff polynomials. This algebraic explosion has limited applications of the linear framework. However, recent results on the graph-based prime factorisation of Kirchhoff polynomials may help subdue the combinatorial complexity. By prime decomposing the graphs contained in an expression of Kirchhoff polynomials and identifying the graphs giving rise to equal polynomials, we formulate a coarse-grained variant of the expression suitable for symbolic simplification. We devise criteria to efficiently test the equality of Kirchhoff polynomials and propose two heuristic algorithms to explicitly generate individual Kirchhoff polynomials in a compressed form; they are inspired by algebraic simplifications but operate on the corresponding graphs. We illustrate the practicality of the developed theory and algorithms for a diverse set of graphs of different sizes and for non-equilibrium gene regulation analyses.

## 1 Introduction

Linear diffusion processes (of information, probabilities, concentrations) on graph models are abundant in science (1; 2; 3). The great significance of Kirchhoff polynomials (4) stems from their role in linking the graph topologies to the symbolic steady-state expressions of such processes. In biology, analyses of linear diffusion processes on graphs, originating from areas as diverse as enzyme kinetics, G-protein-coupled receptors, and gene regulation, have recently been unified under a common mathematical linear framework (5). The linear framework represents a biological system as a labelled directed graph (graph, for short) having molecular states as vertices, state transitions as edges, and transition rate constants as edge labels. The dynamics on the graph have deterministic (linear ODEs) and stochastic (Markov process master equation) interpretations, and define how states (concentrations, respectively probabilities) evolve over time (6). Closed form steady states of such linear framework models (LFMs) always exist and can be symbolically derived from initial conditions and the basis of the kernel of a matrix representation of LFMs, namely the graph Laplacian matrix (5). For systems at thermodynamic equilibrium, the principle of detailed balance dictates “history-independent” equilibrium steady states, for which the basis elements of the kernel can be derived from products of equilibrium constants along any path in the model graph (5). However, a breakdown of detailed balance occurs when systems expend energy, leading to “history-dependent” non-equilibrium steady states of substantially higher algebraic complexity (7). Namely, away from equilibrium a basis element of the kernel of the graph Laplacian matrix becomes a homogeneous multivariate polynomial called the *Kirchhoff polynomial*, which according to Tutte’s Matrix-Tree Theorem (8) can be equivalently obtained by i) symbolically deriving all (*j, j*)-minors of the graph Laplacian matrix and summing them up, and by ii) enumerating all spanning trees in the model graph, multiplying the symbolic labels in each tree, and adding the resulting monomials of all spanning trees.

Departure from equilibrium and the ensuing “history dependence” pose a fundamental challenge – the number of spanning trees and, correspondingly, the size of their Kirchhoff polynomials and symbolic steady-state expressions frequently grows super-exponentially with the size of the graph models (9). Symbolic derivations bring great benefits in understanding non-equilibrium biological phenomena despite this seemingly unmanageable combinatorial explosion. In consequence, there has been a prolific development of software (reviewed in (10)) and of methods to derive steady states and rate equations of biological models that can be classified as falling within the linear framework using, among others, graph theoretical methods (11; 12), systematic determinant expansion (13), and Wang algebra (14). In computer science, advances have also been made to enumerate the set of all spanning trees from which Kirchhoff polynomials are obtained (15; 16). However, all existing exact methods and algorithms suffer from the aforementioned combinatorial explosion and they only offer limited and *ad hoc* manipulation of steady-state expressions. This hinders the in-depth understanding of the role of energy expenditure in biological systems, the extraction of general principles of eukaryotic gene regulation (7) and differential signalling (17), and the analysis of more detailed models that follow from advanced experimental techniques, such as phosphoproteomics (18).

An important step towards taming the combinatorial complexity is the realisation that a model graph *G* can be efficiently decomposed into smaller graphs whose Kirchhoff polynomials are prime factors of the Kirchhoff polynomial of *G* (19). This graph-based polynomial factorisation provides a natural, compact representation that does not directly depend on the number of spanning trees but rather on directed graph connectivity. Here, we exploit the factorisation to further develop theory and algorithms for dissecting and mitigating the seemingly intractable combinatorial complexity. Our approach aims to simplify expressions of Kirchhoff polynomials, bypassing the customary expensive symbolic generation and manipulation by computer algebra systems. Specifically, we consider the prime factors of all Kirchhoff polynomials in an expression as symbolic variables. The resulting coarse-grained expressions allow for symbolic simplification without explicit generation of the polynomials. To explicitly generate the Kirchhoff polynomials, e.g. as is needed for their repeated evaluation, we propose a recursive and an iterative heuristic algorithm inspired by algebraic simplification, but operating on the graphs alone. Applied to a collection of graphs, in particular, graph connectivity aware heuristics prove to be useful in practice, with large compressions and short running times. Further, we extend the sharpness analysis of gene expression in development from (7) and show that away from equilibrium, four binding sites allow for previously unknown qualitative shapes of gene regulation functions. The methods and algorithms are implemented in the Python package *KirchPy* available on https://gitlab.com/csb.ethz/KirchPy.

## 2 Background

### 2.1 Linear framework models (LFMs)

Let us consider the model of Ca^2+^-dependent nuclear translocation of the nuclear factor of activated T cells (NFAT) from (20), which can be expressed in the linear framework. NFAT can be in one of three states – cytoplasmic phosphorylated 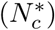, cytoplasmic dephosphorylated (*N*_*c*_), or nuclear (*N*_*n*_), and undergo four reactions – dephosphorylation, phosphorylation, nuclear import, and nuclear export with respective rate constants *r*_1_, *r*_2_, *r*_3_, and *r*_4_. This system functions away from equilibrium because it contains multiple irreversible, and thus energy expending reactions. The reaction scheme of NFAT can formally be represented as a simple labelled directed graph *G* = (*V, E*) (see Figure 1a). The graph *G* is composed of a set of vertices 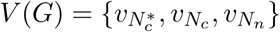 corresponding to NFAT states (correspondence marked in subscript), and a set of edges (ordered pairs of distinct vertices; no multiple parallel edges allowed) 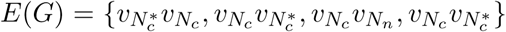 corresponding to reactions. We associate a label *ℓ*(*uv*), standing for a mathematical expression, to each edge *uv* ∈ *E*(*G*). For example, with 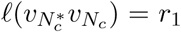 we mark that the label associated to 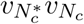 is the rate constant *r*_1_. Additionally, by *ℓ*(*G*) we define the *set of all edge labels* of *G*.

Here, we are primarily interested in LFMs that correspond to *strongly connected* graphs. Namely, graphs *G* in which there exists a directed path from *u* to *v* and from *v* to *u* for any two vertices *u, v* ∈ *V* (*G*) (as in Figure 1a); Figure 1d shows an example for which this property does not hold. However, this does not limit the generality of the developed algorithms and theory.

**Figure 1:**
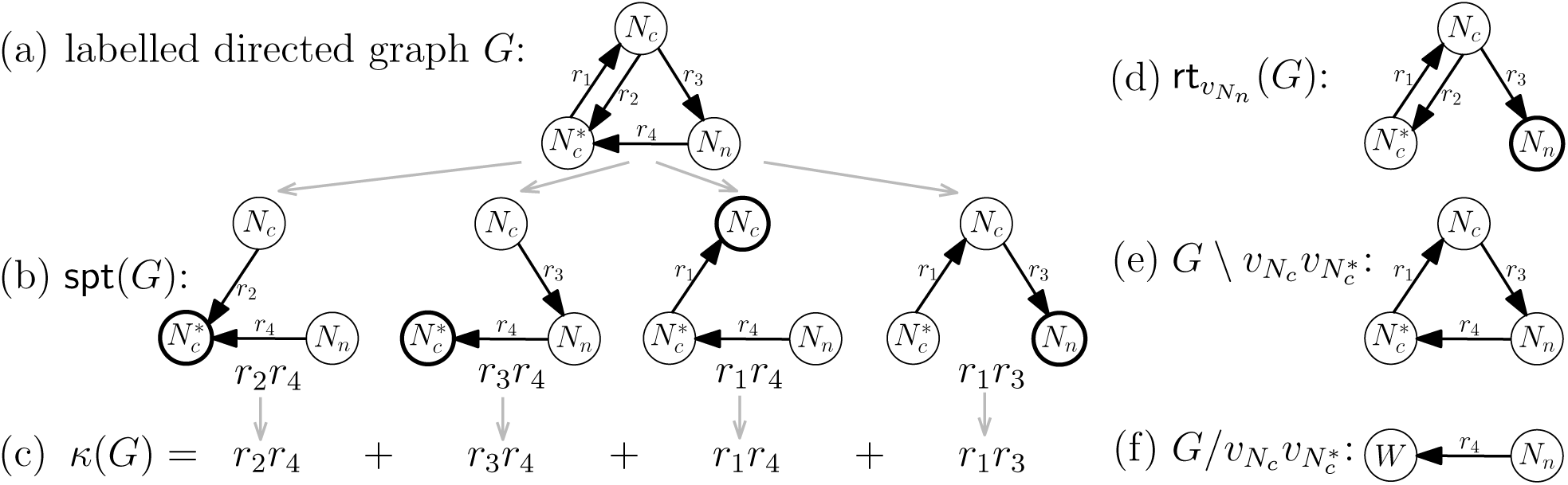
Example model for nuclear translocation of the nuclear factor of activated T cells (NFAT) (20). (a) Graph *G*, (b) all its spanning trees, and (c) the corresponding Kirchhoff polynomial. (d) The graph obtained by rooting *G* at 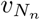, (e) the edge deleted graph 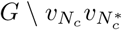, and (f) the edge contracted graph 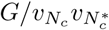. Vertex labels mark the states that they represent and *W* denotes a vertex obtained after edge contraction. Highlighted vertices are, respectively, roots of the corresponding spanning tree and of all spanning trees when rooting a graph.

LFMs are frequently obtained from more complicated models after applying the technique of time-scale separation, stating that a part of a biochemical system operating much faster than the rest of the system can be assumed to have reached a steady state (5). This model reduction could result in edge labels involving (non-linear) algebraic expressions of kinetic parameters and species concentration terms. To retain the linearity of LFMs, concentration terms in labels must correspond to species not contained in *V* (*G*). These could be species acting on the slow time scale or other entities as in the case of NFAT, where *r*_1_ is assumed to be modulated by Ca^2+^ oscillations. We circumvent explicitly dealing with the arbitrary, though biologically significant, algebraic structure of the label expressions by regarding them as uninterpreted symbols *ℓ*(*uv*) that denote unique edge names.

We concentrate on the deterministic interpretation of LFM dynamics (also called Laplacian dynamics) and associate each vertex *v*_*i*_ ∈ *V* (*G*) in *G* to a non-negative species concentration *x*_*i*_ and each edge to a mass-action reaction. In the resulting dynamical system, species concentrations associated to vertices flow in the direction of the edges at rates proportional to the concentrations on the edges’ source vertices, where proportionality is set by the edge label *ℓ*(*uv*).

The example model from Figure 1a is *closed*, it does not exchange matter with the environment. The dynamics of closed LFMs can be expressed in the form:

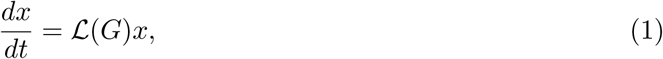

where *x* = (*x*_1_, …, *x*_*n*_)^*T*^ is the vector of species’ concentrations corresponding to each vertex *v*_1_, …, *v*_*n*_ ∈ *V* (*G*) and 𝓛(*G*) is the *graph Laplacian matrix* of *G* defined as:

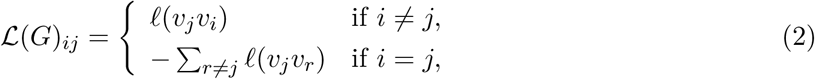

and *ℓ*(*v*_*j*_*v*_*i*_) = 0 when the *v*_*j*_*v*_*i*_ ∉ *E*(*G*). For the example model this means:

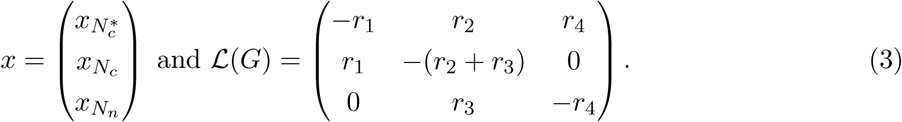

In closed systems the total amount of material *x*_*t*_ is conserved according to a single conservation law *x*_1_ + … + *x*_*n*_ = *x*_*t*_. The system has a unique stable steady state that can be derived symbolically from initial conditions and the kernel of 𝓛(*G*) (6).

Graphs *G* for *open* LFMs with synthesis and degradation reactions are obtained by adding a vertex *v*_Ø_ representing the environment to a *core graph* 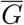 (akin to closed systems, the core graph is composed of all non-synthesis and non-degradation reactions), and by introducing directed edges from *v*_Ø_ to the synthesised species in 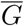 with labels *s*_*i*_ and edges labelled *d*_*i*_ from the degraded species to *v*_Ø_. The dynamics of open LFMs are defined in general form as:

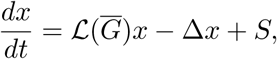

where 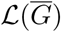 is the graph Laplacian matrix of the core graph, Δ is a diagonal matrix with Δ_*ii*_ = *δ*_*i*_ the degradation rate constants of species *i*, and *S* is a vector *S*_*i*_ = *s*_*i*_ comprising the synthesis rate constants. In open systems, the total amount of matter is not conserved, but synthesis and degradation at steady state are balanced: *δ*_1_*x*_1_ + … + *δ*_*n*_*x*_*n*_ = *s*_1_ + … + *s*_*n*_. Similarly to closed systems, but assuring that the steady state concentration at *v*_Ø_ is always 1, the unique stable steady state for vertex *v*_*i*_ (*v*_*i*_ ≠ *v*_Ø_) can be derived symbolically. For more details on LFMs, see (5; 6; 21).

### 2.2 Spanning trees

A class of *subgraphs*, so-called *spanning trees*, connect non-equilibrium steady states of LFMs to model graph structure.

A graph *H* is a subgraph of a graph *G* if *V* (*H*) ⊆ *V* (*G*) and *E*(*H*) ⊆ *E*(*G*), such that every edge in *G* between vertices in *H* is also an edge in *H*. For *V*′ ⊆ *V* (*G*), *G*[*V*′] denotes the *induced subgraph* of *G* by the set of vertices *V*′. A strongly connected component (*SCC*) of *G* is any largest (w.r.t. vertex inclusion) strongly connected induced subgraph of *G*. The definition implies that no two distinct SCCs share a vertex, that is, the SCCs *G*_1_, …, *G*_*k*_ of a graph *G* induce a unique partition *V* (*G*_1_), …, *V* (*G*_*k*_) of *V* (*G*). Further, two distinct SCCs *G*_*i*_ and *G*_*j*_ can be connected by either a directed path from *G*_*i*_ to *G*_*j*_, or from *G*_*j*_ to *G*_*i*_, but not by both. The existence of such unidirectional paths induces a unique partial order on the SCCs *G*_1_, …, *G*_*k*_.

A *rooted directed spanning tree* (spanning tree, for short) *A* is a subgraph of *G* that spans its vertex set such that there is a unique directed path from any vertex to a root vertex. We denote the set of all spanning trees of *G* by `spt` (*G*), and the set of all spanning trees rooted at a vertex *v* by `spt`_*v*_(*G*) (see Figure 1b for all spanning trees of the example graph). To obtain a graph containing only spanning trees rooted at a vertex *v* we define the graph *rooting* operation `rt`, so that `rt`_*v*_(*G*) is the graph constructed from *G* by removing all edges outgoing from *v* (see Figure 1d). Likewise, we call a graph *G rooted* at a vertex *v* if *v* has no outgoing edges and *v* is reachable from every other vertex in *G*. Observe that `spt`_*v*_(*G*) = `spt` (`rt`_*v*_(*G*)). Graph *G* contains a spanning tree iff the partial order of the SCCs has exactly one maximal element, i.e. no other SCC is reachable from a maximal SCC. Such a maximal SCC is also called a *terminal SCC*.

### 2.3 Kirchhoff polynomials and steady states

A spanning tree *A* of a graph *G* with *n* vertices is a subgraph with *n* − 1 edges *e*_1_, …, *e*_*n*−1_ ∈ *E*(*G*) (we denote edges by *e* when not interested in the pairs of vertices defining them). In a *uniquely labelled graph G*, i.e. when no two edge labels in *G* are the same, *A* can also be represented as a monomial *ℓ*(*e*_1_) *ℓ*(*e*_2_) … *ℓ*(*e*_*n*−1_) in the edge labels of *G*. Further, the set of all spanning trees of *G* can be represented by a homogeneous multivariate polynomial over the variables *ℓ*(*e*_*i*_), *e*_*i*_ ∈ *E*(*G*). This polynomial is called the *Kirchhoff polynomial κ*(*G*) (see Figure 1c):

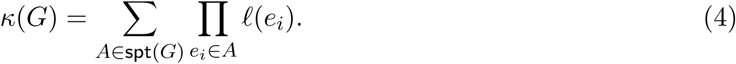

Note that *κ*(*G*) = 0 when no spanning trees exist in *G* and *κ*(*G*) = 1 when *G* consists of a single vertex. We also denote the Kirchhoff polynomial of all spanning trees rooted at vertex *v* by 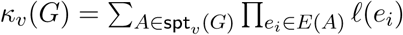 (a shorter notation for *κ*(`rt`_*v*_(*G*))).

A Kirchhoff polynomial *κ*(*G*) can have multiple *algebraically equivalent representations* Γ_*i*_(*κ*(*G*)) (*i* indexes all such representations) corresponding to different *expression trees* (see Figure 2). We consider expression trees in which the branch vertices represent the operations of *n*-ary addition or multiplication, and leaf vertices are the unique edge labels *ℓ*(*G*), the variables of *κ*(*G*). We define the *size* of a representation of *κ*(*G*), |Γ(*κ*(*G*))|, as the size of its corresponding expression tree. However, a change of variables requires an extended definition because it produces a forest of expression trees, and not a single tree (see Figure 2c). We define the size of a Kirchhoff polynomial in such a representation as the total number of branch vertices and leaves in the forest plus the number of expression trees in the forest. We need to account for the number of expression trees since each of them has a unique pointer indicating its location within the other expression trees. The unique steady state of LFMs can be symbolically obtained from initial conditions and the kernel of the graph Laplacian matrix by employing *Tutte’s Matrix-Tree Theorem* (8).

**Figure 2:**
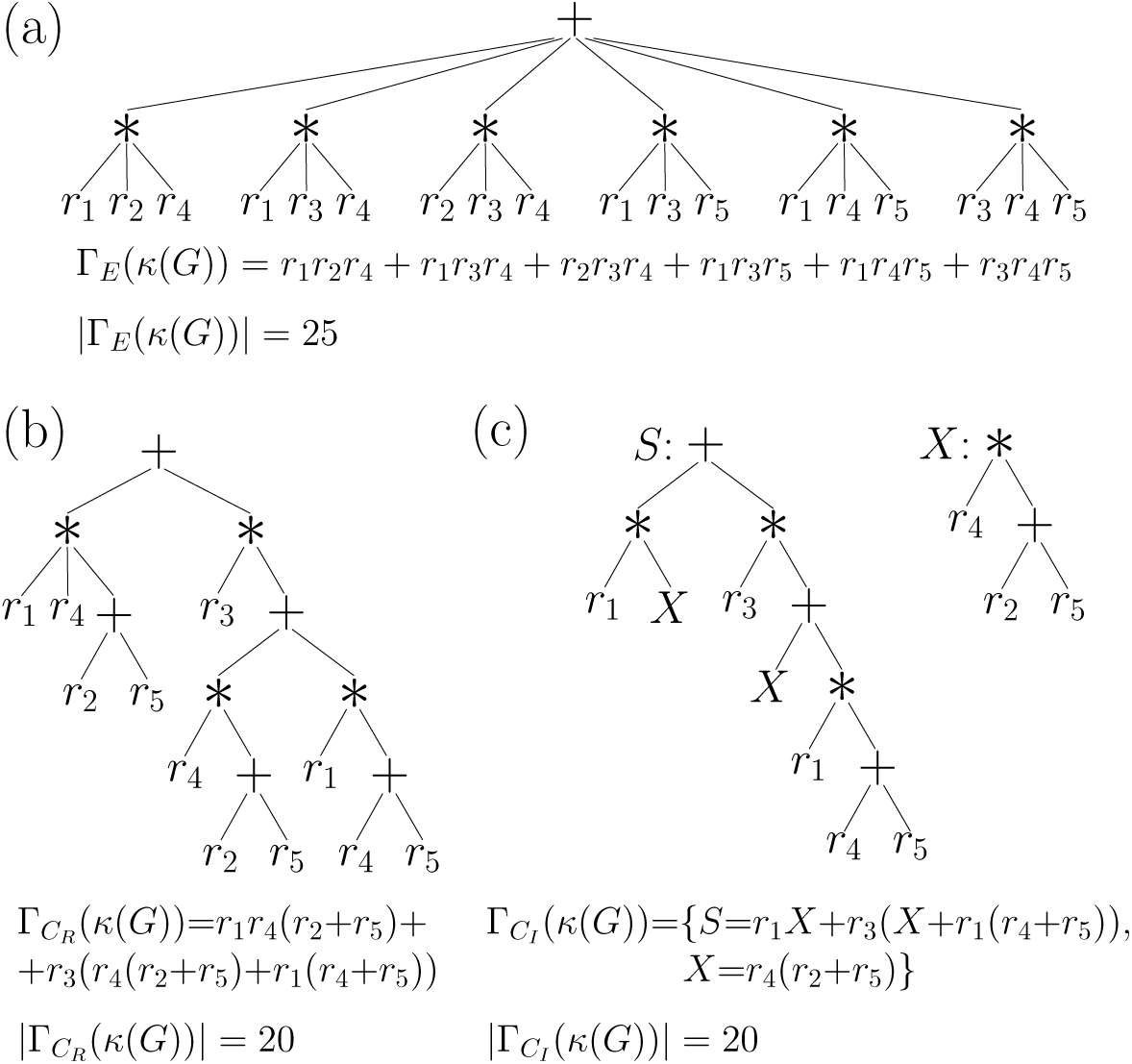
Algebraically equivalent representations of a Kirchhoff polynomial, their expression trees, and sizes for an example graph *G* (see Figure S1) in (a) the fully expanded representation, (b) a simplified representation, e.g. obtained by algorithm *C*_*R*_, and (c) a change of variables form (forest of expression trees), e.g. obtained by algorithm *C*_*I*_. The size of a representation is the sum of the numbers of branch vertices and of leaves in the expression tree. With change of variables, each expression tree from the forest is assigned a pointer counting as 1 to the size of the representation and pointing to the leaves of other expression trees where it should be substituted to obtain the expression tree of the complete Kirchhoff polynomial. The pointer *S* denotes the “starting” tree.

#### Theorem 2.1

(Tutte’s Matrix-Tree Theorem). *Let G be a graph with n vertices then the minors* 𝓛(*G*)_(*i,j*)_ *of its Laplacian matrix can be expressed, up to a sign, by the Kirchhoff polynomial rooted at the vertex v*_*j*_ *corresponding to the j-th column of* 𝓛(*G*) *as:*

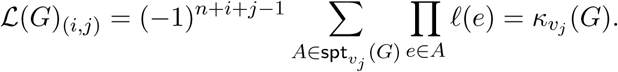

As a result, the non-equilibrium steady-state concentration 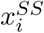 of species *i* associated to vertex *v*_*i*_ in a closed LFM with a strongly connected graph *G* is a fraction of Kirchhoff polynomials:

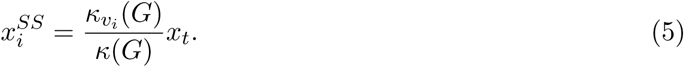

Correspondingly, for open systems and a vertex *v*_*i*_ ≠ *v*_Ø_:

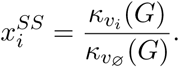

Note that the Kirchhoff polynomial *κ*(*G*) in the denominator of the steady-state expression for closed systems acts as a non-equilibrium partition function (7). For more details on LFMs, derivations, equilibrium steady states, and steady states in non-strongly connected graphs see (5; 6; 21).

### 2.4 Deletion, contraction, prime factorisation

By *G* \ *e* we denote the graph obtained from *G* by deleting edge *e* ∈ *E*(*G*) (see Figure 1e). Additionally, for a graph *G* and an edge *e* = *v*_*i*_*v*_*j*_ ∈ *E*(*G*), *G/e* is the *edge contracted* graph constructed from *G* by (i) removing the edge *v*_*j*_*v*_*i*_, if it exists, and all out-going edges from *v*_*i*_, i.e. *v*_*i*_*u* ∈ *E*(*G*) and (ii) fusing vertices *v*_*i*_ and *v*_*j*_ into a new vertex *w* (see Figure 1f). Edge contractions may give rise to graphs with multiple parallel edges between two vertices. To correct this we replace *m* multiple parallel edges *e*_1_, *e*_2_,…, *e*_*m*_ from *u* to *v* with a single edge *e* = *uv* so that *ℓ*(*e*) = *ℓ*(*e*_1_) + *ℓ*(*e*_2_) + … + *ℓ*(*e*_*m*_).

The defined graph operations can be used to decompose *κ*(*G*), given an edge *e* ∈ *E*(*G*), into a sum of Kirchhoff polynomials according to the classic *deletion-contraction* identity (22):

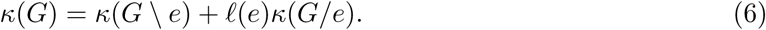

A Kirchhoff polynomial *P* is a *factor* of another Kirchhoff polynomial *Q*, if there exists a Kirchhoff polynomial *R* such that *Q* = *P* · *R*. A Kirchhoff polynomial *P* that cannot be factorized into non-trivial factors is called *prime*. Correspondingly, for graphs, *G*′ *is a component (a prime component) of G* if *κ*(*G*′) *is a factor (a prime factor) of κ*(*G*). Reference (19) introduces graph decomposition rules that correspond to factorisation steps of the Kirchhoff polynomial. In particular, the method yields in linear time graphs whose Kirchhoff polynomials are prime factors of the Kirchhoff polynomial of the original *G*:

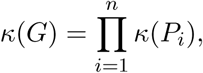

where *P*_*i*_ are the prime components of *G*. A prime component *P*_*i*_ can be either (i) strongly connected or (ii) rooted at *v* such that *P*_*i*_ \ *v* (here \ denotes vertex deletion) is strongly connected and *P*_*i*_ does not have any non-trivial vertex dominators (a vertex *u* dominates a vertex *w* if every path from *w* to *v* goes through *u*). We call graphs with prime Kirchhoff polynomials also *prime graphs*. Importantly, the prime factorisation is conditional on label uniqueness—when different edges have equal labels or there are variables shared across labels, the factorisation is not guaranteed to be prime.

## 3 Efficient manipulation of Kirchhoff polynomials

Non-equilibrium steady states of LFMs are ratios (or more generally: expressions) of Kirchhoff polynomials. Similarly, any symbolic expression derived from steady state LFMs through arithmetic and calculus will also comprise expressions of Kirchhoff polynomials. Examples are ratios of steady states, steady-state rate equations, *EC*_50_ values for steady-state dose-response curves, differential responses (17), and steady-state parameter sensitivities (expressions differentiated with respect to a reaction constant). Correspondingly, algebraic manipulation of expressions of Kirchhoff polynomials is important to understand when expressions can be simplified. For example, the steady state of a LFM can be simplified if the numerator and denominator share common factors. After all common factors are crossed out, numerator and denominator become relatively prime and further simplification is not possible.

To circumvent tedious symbolic manipulation of combinatorially complex algebraic expressions, we exploit properties of Kirchhoff polynomials that allow their implicit manipulation, that is, without explicitly generating polynomials in expanded form but working with the corresponding graphs. More precisely, we (i) find the prime components corresponding to prime factors of all Kirchhoff polynomials in the expression, (ii) determine which prime components generate identical Kirchhoff polynomials, and (iii) form a *coarse-grained representation* of the original expression by substituting prime components with symbolic variables, where prime graphs with equal Kirchhoff polynomials are assigned the same variable, and finally, (iv) symbolically simplify the coarse-grained expression. However, it is an open problem to efficiently determine which prime components generate equal Kirchhoff polynomials without their explicit generation.

### 3.1 Prime graphs with equal Kirchhoff polynomials

We consider Kirchhoff polynomial equality in the algebraic sense. By uniquely labelling a graph we assign identity to each edge through its label, that is, a label defines a particular reaction. Applying the graph operations of prime decomposition, edge deletion, edge contraction, and vertex rooting to a uniquely labelled graph preserves the identity of the reactions while the names of the vertices can change. However, when comparing two Kirchhoff polynomials originating from different sources, identical (different) reactions between the sources need to carry the same (different) labels to have a meaningful comparison.

A necessary condition for two polynomials to be equal is that they have the same set of variables corresponding to terms with non-zero coefficients. This condition cannot be transferred directly to compare the graphs generating Kirchhoff polynomials because the graphs may contain *nuisance edges* that do not participate in any spanning tree. With nuisance edges, the set of labels of two graphs that generate equal Kirchhoff polynomials will be different. However, if we compare only prime graphs we can prove that they do not contain nuisance edges because every edge participates in at least one spanning tree.

#### Theorem 3.1.

*Let G be a prime graph, then each edge in G participates in at least one spanning tree.*

Absence of nuisance edges in prime graphs allows us to formulate a necessary condition for Kirchhoff polynomial equality:

#### Corollary 3.2.

*Let G and H be two prime graphs with equal Kirchhoff polynomials, then G and H have equal sets of edge labels, i.e. κ*(*G*) = *κ*(*H*) ⇒ *ℓ*(*G*) = *ℓ*(*H*).

*Proof.* Follows directly from Theorem 3.1.□

The condition can be tested efficiently since it involves only a comparison between sets, but it is not a sufficient condition for Kirchhoff polynomial equality (see Figure 3a).

**Figure 3:**
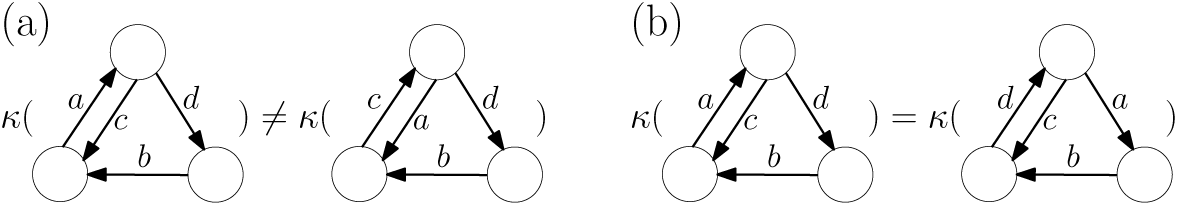
Examples for (a) two graphs with equal edge label sets but different Kirchhoff polynomials and for (b) two non-*λ*-isomorphic prime graphs with identical Kirchhoff polynomials.

To obtain a graph-based sufficient condition of Kirchhoff polynomial equality, we define the term *λ-isomorphism* to denote a vertex bijection that is edge-preserving and enforces the corresponding edges to have identical labels.

#### Definition 3.3

(*λ*-isomorphism). *Two labelled graphs G and H are called λ-isomorphic, denoted G* ≃_*λ*_ *H, iff there exists a bijective mapping ψ* : *V* (*G*) ↦ *V* (*H*), *such that:*

1. *uv* ∈ *E*(*G*) *iff ψ*(*u*)*ψ*(*v*) ∈ *E*(*H*) *and*
2. *ℓ*(*uv*) = *ℓ*(*ψ*(*u*)*ψ*(*v*)).

Evidently, two *λ*-isomorphic graphs give rise to equal Kirchhoff polynomials because the graphs differ only by vertex names, and otherwise have identical topology and labels.

#### Observation 3.4.

*Let G and H be λ-isomorphic, then they generate identical Kirchhoff polynomials, i.e. G* ≃_*λ*_ *H* ⇒ *κ*(*G*) = *κ*(*H*).

To derive a condition testing for *λ*-isomorphism we first define the so-called *line graph* ℒ(*G*) associated to *G*.

#### Definition 3.5

(Line graph). *The line graph* ℒ(*G*) *associated to the graph G satisfies the conditions:*

1. *the vertices of* ℒ(*G*) *are the* unique edge labels *of G, i.e. V* (ℒ(*G*)) ≡*ℓ*(*G*) *and*
2. *two vertices u, v* ∈ *V* (ℒ(*G*)) *are joined by an edge uv iff u* = *ℓ*(*rs*), *v* = *ℓ*(*st*) *for r, s, t* ∈ *V* (*G*).

#### Theorem 3.6.

*Two prime graphs G and H are λ-isomorphic iff the edge sets of their line graphs are equal, i.e. G* ≃_*λ*_ *H* ⇔ *E*(ℒ(*G*)) = *E*(ℒ(*H*)).

Theorem 3.6 allows us to formulate a sufficient condition for prime Kirchhoff polynomial equality:

#### Corollary 3.7.

*Let G and H be two uniquely labelled prime graphs whose line graphs have equal edge sets, then the Kirchhoff polynomials they generate are equal, i.e. E*(ℒ(*G*)) = *E*(ℒ(*H*)) ⇒ *κ*(*G*) = *κ*(*H*).

*Proof.* Follows directly from Observation 3.4 and Theorem 3.6. □

The sufficient condition in Corollary 3.7 is also cheap to evaluate since it only involves line graph construction, which has quadratic time complexity, and the comparison of two sets. The condition is not necessary for prime Kirchhoff polynomial equality (see Figure 3b).

### 3.2 Formulation of coarse-grained expressions

We use the conditions in Corollary 3.2 and Corollary 3.7 to assign identical variable names to prime graphs with equal Kirchhoff polynomials when formulating the coarse-grained description of an expression of Kirchhoff polynomials without their explicit generation. First we apply the necessary condition to filter possible matches, and then the sufficient one to certify the equality.

Pairs of prime graphs that are non-*λ*-isomorphic but have the same label sets require special attention. With such pairs, we cannot guarantee that we have identified all graphs with equal Kirchhoff polynomials, which translates to a lack of guarantees for maximal symbolic simplification of the coarse-grained description. However, without such pairs of graphs in the expression, we can guarantee the exhaustive identification of prime graphs with equal Kirchhoff polynomials. Further, comparisons of prime graphs can be accelerated by realizing that each prime component of a graph is equal to at most one prime component of another graph, since prime factorisation partitions the set of labels. Note that there might be other reasons that do not guarantee full simplification and contexts in which full simplification is guaranteed (see Supplementary Material for details). Additionally, some proofs and derivations assume that the graph models have unique and irreducible expressions in their labels. If this assumption is not met, e.g. when different reactions have the same rate constant, rate constants are expressions that can be simplified, or rate constants contain symbols shared across different labels, then additional symbolic simplification might be required since the primality of the decomposition is not guaranteed and the manipulation formulas of the coarse-grained representation might not hold.

The study of non-equilibrium steady-state LFMs is not complete without efficient methods to apply differentiation and integration to Kirchhoff polynomials, for example, to derive parameter sensitivities. In Supplementary Material we show that the properties of Kirchhoff polynomials allow us to map differentiation and integration to graph operations, and thus to work with the implicit coarse-grained representation.

### 3.3 Application examples

To illustrate the manipulation of expressions of Kirchhoff polynomials in the coarse-grained representation, we first consider a simple open receptor trafficking model with graph *G* shown in Figure 4a. It consists of species for an unbound surface receptor *R*, a cell surface ligand-receptor complex *RL*, their respective internalised counterparts *R*_*i*_ and *RL*_*i*_, and a set of state transition, synthesis, and degradation reactions. Figure 4b shows the steady state for *RL*_*i*_ obtained by prime decomposing the graphs in the steady-state ratio and crossing out the common factors. Without complete generation of the polynomials *κ*_*RL*_*i* (*G*) or *κ*_Ø_(*G*), we immediately see that the resulting expression does not depend on the rate constants *r*_1_, *r*_2_, and *r*_3_.

**Figure 4:**
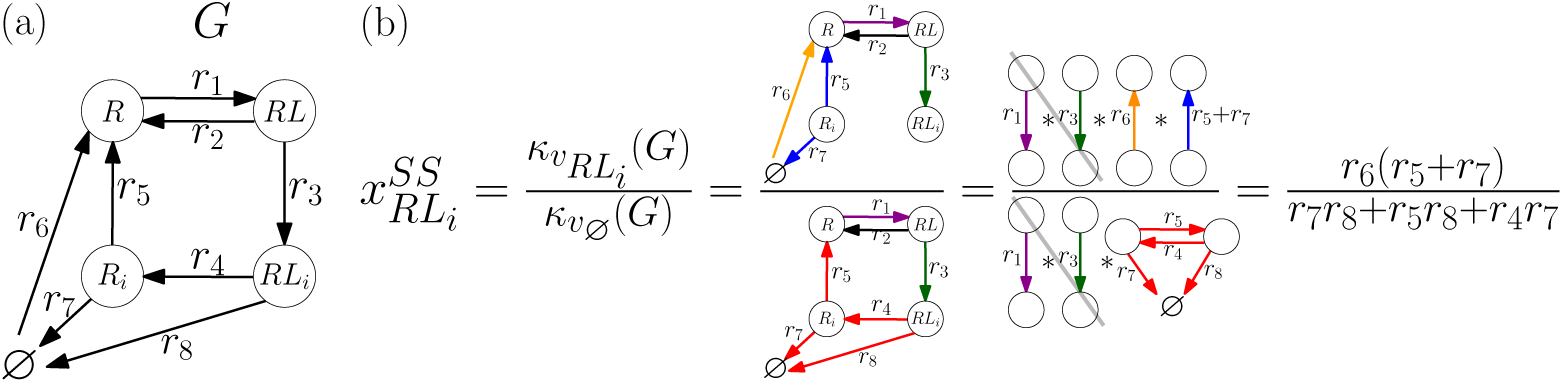
Simplification of expressions of Kirchhoff polynomials in the coarse-grained representation. (a) Simple receptor trafficking model *G* and (b) simplification of its steady-state expression. Prime components with equal Kirchhoff polynomials have the same colour; reaction constant *r*_2_ is a label on a nuisance edge marked in black and does not partake in any prime component. Note that vertex labels are not important since they can change during edge contractions. Also, the symbol *κ* is omitted in front of the graphs for clarity.

The coarse-grained description is most instrumental in understanding large non-equilibrium systems when steady-state derivations are difficult or practically impossible. For example, one could study relative responses, expressed through ratios between the steady states of two species, which upon simplification could become decoupled from a subset of reaction rate constants. Biologically important examples for such relative responses are ratios of folded and misfolded protein conformations in proteostasis, where chaperones expend energy to alter the ratios (23), and ratiometric mechanisms in signalling, in which the ratio between unoccupied receptors and ligand-bound receptor complexes determines downstream effects (24). Decoupling can be used to infer the connectivity of reaction networks, or design measurements to focus on (or isolate) the effect of certain reactions on the combinatorially complex steady state of a system.

Let us consider the detailed catalytic cycle of the Prostaglandin H Synthase 1 (PGHS) from (25) (for description see COX in Table S1), whose graph *G* is shown in Figure S2a and contains 24 quadrillion spanning trees. We analyse the decoupling from reaction rate constants in all of its possible steady-state ratios by coarse-graining the relevant Kirchhoff polynomials and cancelling the common factors between the numerator and denominator. We find that the ratio between the steady states of species *E*21 and *E*17 (two states in the peroxidase cycle of the enzyme containing the arachidonic acid radical in the cyclooxygenase site and differing by the state of Tyrosine 385) contains the fewest number of reaction dependencies – only five (see Figure S2b) – while the Kirchhoff polynomials *κ*_*E*21_(*G*) and *κ*_*E*17_(*G*) consist of trillions of spanning trees.

## 4 Compact generation of Kirchhoff polynomials

After simplifying an expression of Kirchhoff polynomials in its coarse-grained form, we find which labels have vanished (the corresponding reactions do not affect the expression) and which ones remain (the corresponding reactions might affect the function modelled by the expression). However, to symbolically obtain the simplified expression for further analysis or repeated evaluation, e.g. for parameter space exploration, we have to explicitly generate full-length Kirchhoff polynomials. The coarse-grained representation is advantageous here as well because we only need to generate the Kirchhoff polynomials for prime graphs with unequal Kirchhoff polynomials.

### 4.1 Recursive and iterative algorithms

Specifically, we extend the approach of (19) to Kirchhoff polynomial generation, namely, that of algebraic simplification—compression of the polynomial to an equivalent but more compact form. Thus we look for an algorithm *C* that takes a graph *G* as input and produces a representation of its Kirchhoff polynomial Γ_*C*_ (*κ*(*G*)) of size as small as possible. An ideal algorithm *C* would generate Γ_*C*_ (*κ*(*G*)) in a maximally compact form, bypassing explicit generation and tedious simplification. However, it is hard to even check if a Kirchhoff polynomial is fully simplified. Therefore, we aim to propose algorithms that, without guarantees for maximal compression, provide satisfactory results to practical problems.

The prime decomposition in (19) behaves as the ideal algorithm *C* for compression—in linear time, it produces a guaranteed maximally compact representation for a graph due to the irreducibility of each prime component. However, it cannot be applied to prime graphs, which can also have sizable Kirchhoff polynomials. To compress the Kirchhoff polynomials of the prime components, we rearrange prime Kirchhoff polynomials, particularly by taking a factor out from part of their monomials, such that we can further factorise parts of them. Without explicit generation, this is achieved through the deletion-contraction identity (Equation 6), in which the modified graphs *G* \ *e* and *G/e* could be amenable to further prime decomposition since *e*’s deletion and contraction could change the connectivity of *G*.

With this insight, we formulate the algorithm *C*_*R*_ (initially presented in (19); see pseudocode in Supplementary Material as Algorithm 1). It takes a graph *G*, and recursively alternates between prime decomposition and edge deletion-contraction in every prime component until graphs are reduced to a single vertex or a single edge, whose polynomials are trivial to generate. *C*_*R*_ is easy to implement and produces an expression tree as in Figure 2b that is more compact than the expanded form of the Kirchhoff polynomial. However, multiple recursive calls could unnecessarily work on large graphs with equal Kirchhoff polynomials.

We propose a second, iterative algorithm, *C*_*I*_ (for details and pseudocode see Supplementary Material, Algorithm 2). It employs the graph comparisons certifying Kirchhoff polynomial equality to eliminate the potential redundancy of multiply generating equal Kirchhoff polynomials. In contrast to *C*_*R*_, *C*_*I*_ associates a unique pointer to every graph under study, and reduced graphs are added to a queue for further reduction, while remembering the partial expression tree they participate in. Then, the algorithm iterates over the graphs in the queue, reducing them only if their Kirchhoff polynomials are distinct from the Kirchhoff polynomials of all graphs already considered. Algebraically, this is equivalent to a change of variables—substituting identical parts of the Kirchhoff polynomial with identical symbols and explicitly generating them only once (see Figure 2c for an example). The partial expression trees are then assembled to obtain a forest of expression trees marked with the pointers of the initializing graphs as in Figure 2c. This forest corresponds to a set of Kirchhoff polynomials, which after being substituted into each other, gives rise to the complete Kirchhoff polynomial of graph *G*.

*C*_*I*_’s representation of the Kirchhoff polynomial is more compact than the expanded form and can still be easily evaluated and analysed. However, if there are few small graphs with equal Kirchhoff polynomials encountered during the reduction, compared to *C*_*R*_, *C*_*I*_ might consume more memory (to remember pointers and already considered graphs), have longer running time (due to equality comparisons), and not provide significantly better compression (compare Figure 2b and c). On the other hand, if the reduction encounters many large graphs with equal Kirchhoff polynomials, only *C*_*I*_ may generate practically relevant Kirchhoff polynomials.

An important ingredient of both algorithms *G*_*R*_ and *C*_*I*_ is the choice of an edge for the deletion-contraction operation (function GetEdgeForDelContr in Supplementary Material, Algorithms 1 and 2). It is unknown which edges to delete-contract to generate a maximally compressed Kirchhoff polynomial (19). Therefore, we resort to a heuristic approach: we greedily select an edge to delete-contract such that a criterion on the decomposition properties is optimised. Since Kirchhoff polynomial generation results are instance specific, we explore different heuristics (see Supplementary Material).

### 4.2 Performance evaluation

For performance analysis, we evaluated the running time and compression of *C*_*R*_ and *C*_*I*_, where we define *compression* as the ratio of the size of the expanded representation of a Kirchhoff polynomial |Γ_*E*_(*κ*(*G*))| and the size of its representation produced by an algorithm *C*, |Γ_*C*_(*κ*(*G*))|. Specifically, we applied 109 heuristics on a collection of example graphs of widely different complexity (see Table S1). Ten less complex graph models have tens to millions of spanning trees and two more complex models, HC4 and COXD, have up to quadrillion of spanning trees.

First, we analysed the less complex examples to compare the performance of the different heuristics (see Supplementary Material for details). For more complex graphs, a random heuristic’s performance quickly deteriorates, becoming orders of magnitude worse than heuristics informed by the graph connectivity (Figure S5). We normalized the performance measures over all heuristics separately for each example and divided them into groups (see Supplementary Material, Figures S3 and S4). Post-hoc comparisons of sub-heuristic choices revealed that focusing deletion-contraction on edges relevant to the cycle structure of the graphs, and considering the edge deleted graphs and strongly connected components leads to significantly shorter running time and larger compression on average (see Table S2 and Table S3).

Figure 5 compares the performance of algorithms *C*_*R*_ and *C*_*I*_ for the connectivity-informed heuristics. The performance data for examples of low complexity lie on or symmetrically around the 45°line, indicating that the two algorithms perform alike. However, the more complex the examples, the more apparent becomes the superiority of algorithm *C*_*I*_ over *C*_*R*_. The difference is most striking for compression, implying that the change of variables benefits the compression of all models. Finally, the results for the heuristics leading to the largest compression with *C*_*I*_ (Table 1) show that, for larger graphs, the compressed form is orders of magnitude shorter than the number of spanning trees. Therefore, algebraic compressibility, rather than the number of spanning trees, is a hard bound for Kirchhoff polynomial generation. It is an open problem how to determine the compressibility of a graph, but we can get an impression by comparing HC4 and COXD. The compression results are expected because it is difficult to uncover strong connectivity and domination during the graph reduction procedure in dense graphs with many reversible edges; it is simpler to break open cycles in graphs with low density and many unidirectional edges.

**Table 1:** Performance of algorithm *C*_*I*_ with the heuristics leading to the largest compression on a set of example graphs (see graph descriptions in Table S1): size of the expression tree of the expanded Kirchhoff polynomial |Γ_*E*_(*κ*(*G*))|, size of the compressed expression tree |Γ_*C*_ (*κ*(*G*))|, heuristic 𝓗, compression calculated as the ratio 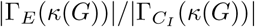, and average running time in seconds obtained from 10 runs (one run for HC4 and COXD) of KirchPy on a Dell laptop with Intel i-7 CPU@2.10GHz and 8GB RAM.

| $G$ | $\vert \Gamma_E(\kappa(G))\vert $ | $\vert \Gamma_{C_I}(\kappa(G))\vert $ | $\mathcal{H}$ | Compression | Time (s) |
| --- | --- | --- | --- | --- | --- |
| COLE1 | 157 | 46 | 2012 | 3.4 | 0.09 |
| AMPAR | 211 | 63 | 2102 | 3.3 | 0.07 |
| MDH | 1,270 | 141 | 2001 | 9.0 | 0.27 |
| ACTMYO | 3,561 | 142 | 3003 | 25.1 | 0.19 |
| KNF33 | 15,553 | 940 | 3114 | 16.5 | 1.41 |
| SHPIL | 45,601 | 786 | 2204 | 58.0 | 1.58 |
| GR | 65,742 | 1,280 | 2102 | 51.4 | 1.63 |
| PHO5 | 640,513 | 4,691 | 3215 | 136.5 | 10.61 |
| RND | 967,681 | 3,281 | 1011 | 294.9 | 7.53 |
| TF | 38,746,801 | 1,191 | 2002 | 32,533 | 1.84 |
| HC4 | 679,477,249 | 333,599 | 1001 | 2,036 | 1,797 |
| COXD | 367,647,474,<br>647,060,221 | 89,532 | 1010 | $4.1 \cdot 10^{12}$ | 661 |

**Figure 5:**
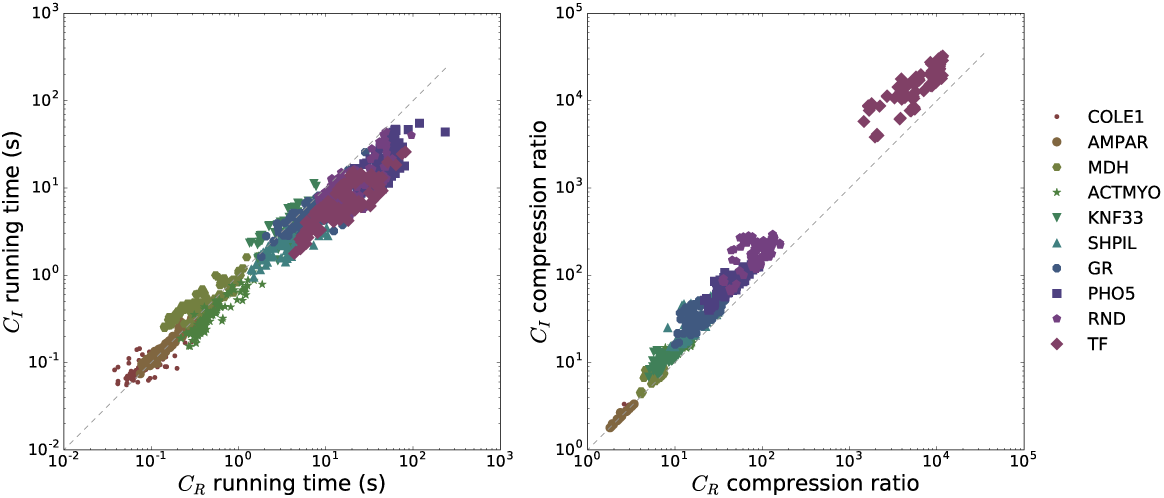
Running time (left) and compression (right) of algorithms *C*_*R*_ and *C*_*I*_ on the collection of less complex examples in Table S1. Each point represents the running time/compression for an example graph obtained by *C*_*R*_ and *C*_*I*_ using the same connectivity-informed heuristic. The dashed line marks equal performance for *C*_*R*_ and *C*_*I*_. Examples are sorted by complexity (number of spanning trees): COLE1 has the lowest complexity and TF the highest.

### 4.3 Application to non-equilibrium gene regulation

Example HC4 from Table 1 belongs to a family of LFMs used in (7) to explore possible biophysical mechanisms behind the sharp expression profile of the *hunchback* gene as a function of the transcription factor Bicoid in the early *Drosophila* embryo. In this family of hypercube graphs, vertices represent DNA microstates (patterns of a transcription factor (TF) bound to a gene), and edges and edge labels mark TF binding (with rates dependent on TF concentration) and unbinding. Graph topology is determined by the number *n* of TF binding sites at the gene, for example, *n* = 3 corresponds to a cube graph (see Figure S6) and *n* = 4 to the four dimensional hypercube HC4. Under the stochastic interpretation of LFM dynamics, microstate probabilities evolve depending on the transition rates until a steady state is reached.

Reference (7) develops a sharpness analysis for relations between gene expression rate and TF concentration, called gene regulation functions (GRFs), that are derived from these LFMs. More precisely, GRFs are functions of steady-state microstate probabilities determined by a choice of an expression strategy. For example, in the *all-or-nothing* strategy, transcription is proportional to the steady-state probability of the microstate in which all TF sites are bound; microstate probabilities are, in turn, functions of TF concentrations. After normalisation, two features are extracted from a GRF to evaluate its sharpness: i) *steepness* – the GRF’s maximal derivative and ii) *position* – the TF concentration at which the maximal derivative is attained. A subsequent exploration of the GRF parameter space by a biased sampling algorithm aims to determine the boundaries of the feasible position-steepness region.

An important result of (7) is that energy expenditure is one possible explanation for the observed sharp response expression profiles in development. In particular, at thermodynamic equilibrium GRF position-steepness regions are restricted by the Hill function, which acts as a *Hopfield barrier*, whereas energy expenditure broadens the feasible position-steepness regions and allows for sharper responses. However, due to the large algebraic complexity of non-equilibrium steady states, the non-equilibrium position-steepness analysis in (7) is limited to models with up to *n* = 3 sites, while the *hunchback* P2 enhancer has 5–7 Bicoid binding sites.

The 2000-fold compression of the *n* = 4 sites model HC4 allows us to extend the non-equilibrium case analysis and explore how the number of binding sites affects the position-steepness regions. We focus on the all-or-nothing expression strategy and models with *n* = 2, 3, 4 binding sites whose non-dimensionalised kinetic parameters are sampled in the range [10^3^, 10^−3^]. We obtain position-steepness boundaries as described in (7), with differences mentioned in Supplementary Material. Our results are shown in Figure 6 and indicate that the *n* = 3 and *n* = 4 site models can achieve steepness of around 5.5 and 6.4, respectively, both exceeding the experimentally fitted Hill coefficient of 5 to the *hunchback* expression profile in response to Bicoid. Interestingly, *n* = 4 site models can generate GRFs with position values greater than 1, which are not attainable by Hill functions, equilibrium GRFs (since they are bounded by Hill functions acting as a Hopfield barrier), and non-equilibrium models with a lower number of sites. A normalised position value of 1 corresponds to a TF concentration at which the GRF is half-maximal, suggesting that *n* ≥ 4 binding sites permit a wider class of GRF shapes in which the maximal steepness arises at TF concentrations larger or equal to the half-maximal TF concentration (see examples in Figure S7). Note that the increase of the position-steepness area from *n* = 2 to *n* = 3 sites is much larger than from *n* = 3 to *n* = 4 sites. We believe that, because of the vastly different dimensions of the sampled parameter spaces (6, 22, and 62 free parameters for models with *n* = 2, 3, and 4 sites, respectively), the more the sites, the less precise the boundaries.

**Figure 6:**
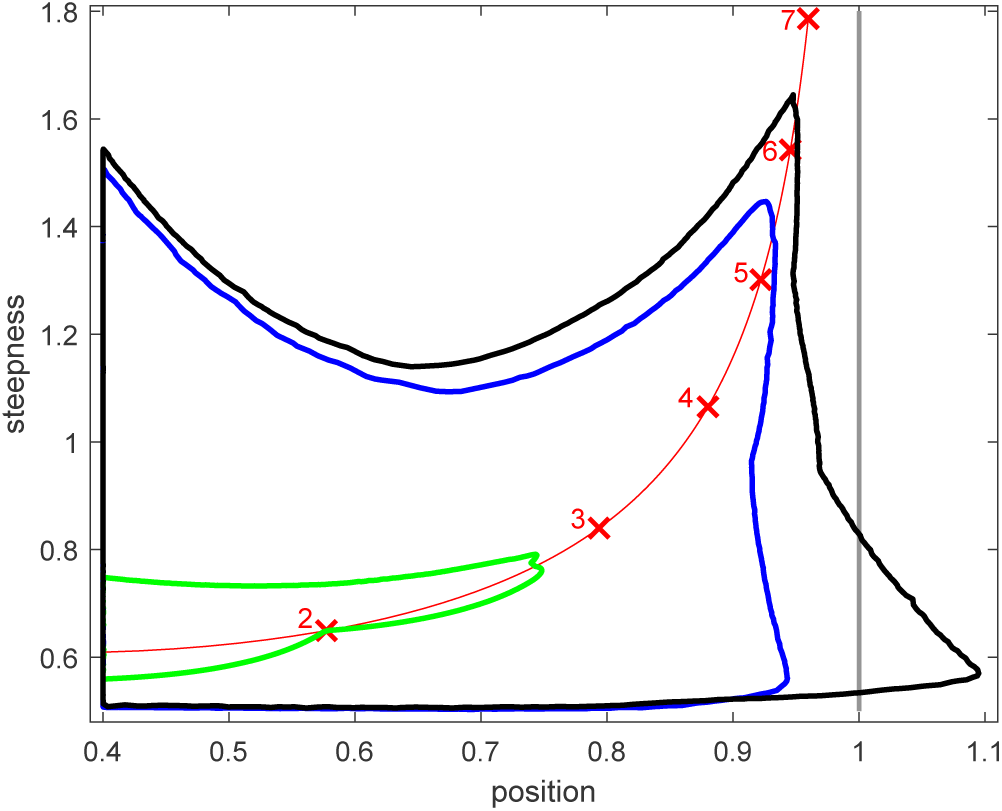
Position-steepness regions for non-equilibrium gene response functions corresponding to models with *n* = 2 (green), *n* = 3 (blue), *n* = 4 (black) transcription factor binding sites. The boundaries are obtained for the all-or-nothing expression strategy by sampling parameter values in the interval [10^3^, 10^−3^]. The *Hill line* defined by the position-steepness loci of Hill functions with coefficients ranging from 1.5 to 7 is shown in red; loci corresponding to integer Hill coefficients are marked with red crosses and numbers. The position asymptote of the Hill line is marked in grey.

## 5 Conclusions

Here, we concentrated on biochemical models falling under the linear framework (5) and took an algebraic graph theory approach to describe their non-equilibrium steady states. The convenient correspondence between graphs and polynomials was essential in the development of theory and algorithms allowing us to manipulate and compress the combinatorially complex expressions. In particular, our coarse-grained representation permits the manipulation of otherwise symbolically intractable expressions of Kirchhoff polynomials. It also helps establish a structure-function relationship between a model and its steady-state response by identifying which reactions do not partake in the expression due to algebraic simplification and, in some cases, which reactions participate in it due to irreducibility.

To explicitly generate individual Kirchhoff polynomials from a simplified coarse-grained expression, our two proposed algorithms produce compressed polynomials that are algebraically equivalent to their fully expanded counterparts. We demonstrated the practical utility of the algorithms for a wide range of graph examples. The large compression results affirm the finding from (19) that Kirchhoff polynomial generation depends on graph connectivity and not on the (super) exponentially growing number of spanning trees. This calls for a more in-depth characterisation of Kirchhoff polynomial compressibility based on connectivity. Additionally, compression allowed us to expand the non-equilibrium gene regulation analysis of (7) and conclude that with four transcription factor binding sites qualitatively different shapes of gene regulation functions can be obtained.

The presented manipulation and generation theory and algorithms are implemented in the Python package KirchPy (available on https://gitlab.com/csb.ethz/KirchPy). A direction for improvement of the manipulation and simplification tools is the further development of Kirchhoff polynomial equality conditions, since the ones we present are not simultaneously necessary and sufficient, such that we cannot, in general, guarantee to identify all graphs giving rise to equal Kirchhoff polynomials in a coarse-grained expression. We anticipate that developments in Kirchhoff polynomial isomorphism similar to those for undirected graphs (26) and optimized deletion-contraction heuristics fuelled by recent advances in strong connectivity and 2-connectivity (27) could further improve the performance of our compression algorithms. Overall, we believe that KirchPy, together with the theoretical insights into manipulation and generation of Kirchhoff polynomials, would i) allow modelling and analysis efforts to catch up with the ever more comprehensive experimental data by promoting the construction and analysis of larger linear framework models, ii) enable the analysis of the functional significance of simplifications in classes of models, which can be useful in experimental design as well as for studying phenomena such as proteostasis, ratiometric and differential signalling, iii) find applications beyond biology because of the equivalence of linear framework models to continuous time Markov Chains, and iv) offer an alternative for steady state derivations that is more convenient than the direct application of the Matrix-Tree Theorem through naive spanning tree enumeration.

## Acknowledgements

We are grateful to Przemyslaw Uznański for insightful discussions and to Jeremy Gunawardena for critical comments and valuable suggestions.

## Funding

This work has been supported by the EU FP7 project IFNAction (contract 223608).

## Author contributions

Conceptualization, P.Y. and J.S.; Methodology and algorithms, P.Y.; Investigation, P.Y.; Writing, P.Y. and J.S. All authors gave final approval for publication.

## Competing interests

We have no competing interests.

## Notes

https://gitlab.com/csb.ethz/kirchpy

