## Supplementary Material for "Efficient Manipulation and Generation of Kirchhoff Polynomials for the Analysis of Non-equilibrium Biochemical Reaction Networks"

#### 1 Prime graphs with equal Kirchhoff polynomials – omitted proofs

Note that we consider in-directed spanning trees as defined in the main text since they are relevant to derive steady states of linear framework models and not out-directed spanning trees, i.e. rooted directed spanning trees for which all edges are directed away from a root vertex, as in (1). Additionally, in (1) rooted directed spanning trees are synonymously termed arborescences. There is a bijection between the in-directed spanning trees of a graph $G$ and the out-directed spanning trees of the reverse (with all edges reversed) graph of $G$. Thus statements on out-directed spanning trees can trivially be rewritten for in-directed ones and vice versa.

**Theorem 3.1.** *Let $G$ be a prime graph, then each edge in $G$ participates in at least one spanning tree.*

*Proof.* The theorem statement is equivalent to the statement that there are no nuisance edges, or formally $\nexists e \in E(G)$ such that $\mathsf{spt}(G/e) = \emptyset$. Assume the contrary, i.e. let $e = uv$ be such that $\mathsf{spt}(G/e) = \emptyset$. Such a graph $G/e$ with no spanning trees is either disconnected or has more than one terminal SCC. Let us investigate the two classes of prime graphs from (1):

- **$G$ is strongly connected.**
  Normal edge contraction, i.e. fusing $u$ and $v$ to obtain a new vertex $u^*$, cannot disconnect or introduce a terminal SCC in a strongly connected $G$. However, the definition of edge contraction w.r.t. spanning trees requires the deletion of edges out-going from $u$ in $G$, which we call $uw_i$. Deleting $uw_i$ upon contraction of $uv$ cannot disconnect $G$ since there exists a directed path from $w_i$ to $u$ due to strong connectivity whose existence is not affected by the contraction. Deleting $uw_i$ also cannot lead to more than one terminal SCC but only multiple new initial SCCs ($w_i$ always has an out-degree of at least one and could become part of a new initial SCC when $uw_i$ is its only incoming edge; $u^*$ can become part of a single terminal SCC). Therefore, $\mathsf{spt}(G/e) \neq \emptyset$.

- **$G$ is a graph rooted at $v_{aux}$, such that $G \setminus \{v_{aux}\}$ is strongly connected and $G$ has no non-trivial dominators.**
  Notice that the edges $u_i v_{aux}$, $u_i \in V(G) \setminus \{v_{aux}\}$, always participate in at least one spanning tree of $G$ since there exists at least one spanning tree $A$ rooted at $u_i$ in the strongly connected $G \setminus \{v_{aux}\}$ and, further, adding $u_i v_{aux}$ to $A$ produces a spanning tree of $G$. Also, notice that in this class of prime graphs (apart from the trivial case when the graph is a single edge) at least two different vertices, e.g. $u_1, u_2 \in V(G) \setminus \{v_{aux}\}$, form an edge with $v_{aux}$. If there is a single edge containing $v_{aux}$, e.g. $u_1 v_{aux}$, $u_1$ would dominate $V(G) \setminus \{v_{aux}, u_1\}$ and thus $G$ has non-trivial dominators.Therefore, $e$ cannot be any of $u_i v_{aux}$.

Thus let $e = uv(u, v \in V(G) \setminus \{v_{aux}\})$. Again, this means that contracting $uv$ either disconnects $G$ or creates at least one additional terminal SCC in it (by definition the only terminal SCC is $v_{aux}$). Contraction of $uv$ deletes out-going edges from $u$, which we again call $uw_i$.

By deleting $uw_i$ ($w_i$ could also be $v_{aux}$) we cannot disconnect $G$ since, first, there exists a path from $w_i$ to $u$ in the strongly connected $G \setminus \{v_{aux}\}$ and, second, there exists a path from $v$ to $v_{aux}$ not passing through $u$ (and thus also from the fused $u^*$ to $v_{aux}$ after the edge deletions induced by the edge contraction are applied) due to the lack of non-trivial dominators. Deleting $uw_i$ also cannot create more than one terminal SCC but only initial SCCs. As before, $w_i$ could become part of new initial SCCs and there could be at most one terminal SCC due to the existence of a directed path from $v$ to $v_{aux}$ not passing through $u$. Therefore, $\mathsf{spt}(G/e) \neq \emptyset$.

□

**Theorem 3.6.** *Two prime graphs $G$ and $H$ are $\lambda$-isomorphic iff the edge sets of their line graphs are equal, i.e. $G \simeq_\lambda H \Leftrightarrow E(\mathscr{L}(G)) = E(\mathscr{L}(H))$.*

*Proof.* The proof follows from (2), Theorem 2, which states that there is a one-to-one correspondence between the set of all graphs with at most one vertex of out-degree zero and at most one vertex of in-degree zero and no isolated vertices, and the set of all line graphs. Prime graphs are either strongly connected or have a single vertex of in-degree zero when considering out-directed spanning trees (and a single vertex of out-degree zero for in-directed spanning trees). Therefore, there is a one-to-one correspondence between prime graphs and the set of all line graphs. This means $\lambda$-isomorphic prime graphs bijectively map to the same line graph (defined by the set of its edges). □

### 2 Limitations of Kirchhoff polynomial coarse-graining

The conditions from Corollary 3.2 and Corollary 3.7 are not necessary and sufficient in general, but they could be such in certain cases. For example, when we compare the Kirchhoff polynomials of two prime graphs $G_1$ and $G_2$ obtained from the same graph $G$ by means of edge deletion and prime decomposition we can be sure that they have the same Kirchhoff polynomials iff they have equal edge label sets. This is because starting from a common uniquely labelled graph $G$, edge deletions and prime decomposition (partition of edges) do not change the comparative topology of $G_1$ and $G_2$. An example for such expressions derived from a single graph by means of edge deletion and prime decomposition are the steady-state expressions of open LFMs – their numerator and denominator contain Kirchhoff polynomials of rooted graphs which could possibly be prime decomposable. Edge contractions, however, have the capacity to permute the edges and lead to the examples in Figure 3.

Note that simplification on a lower or higher level from the coarse-graining could be possible, which prevents full simplification of an expression of Kirchhoff polynomials. One such instance is the difference between two prime Kirchhoff polynomials which share monomials (the corresponding graphs share spanning trees). The common monomials will cancel out if the Kirchhoff polynomials are written in explicit form but cannot be cancelled in the coarse-grained representation. Partially developing the prime Kirchhoff polynomials, e.g. using the edge deletion-contraction identity, to sums containing the shared monomials could often turn to be lengthy. Another instance is when the sum or difference can be reduced by combining Kirchhoff polynomials to form the Kirchhoff polynomial of another graph. For example, according to the edge deletion-contraction identity if we identify graphs $G \setminus e$ and $G/e$ in the expression $\kappa(G \setminus e) + \ell(e)\kappa(G/e)$ we can simplify it to $\kappa(G)$. Other variations of this identity are also possible, e.g. the expression $\frac{\kappa(G)-\kappa(G\setminus e)}{\ell(e)}$ can be written simply as $\kappa(G/e)$.

### 3 Calculus of Kirchhoff polynomials

Taking the partial derivative of a Kirchhoff polynomial with respect to a reaction rate constant $\ell(e)$ corresponding to the label of edge $e$ is equivalent to edge contraction in the corresponding graph as shown by Identity 3.1.

**Identity 3.1.**

$$\frac{\partial \kappa(G)}{\partial \ell(e)} = \kappa(G/e).$$

*Derivation.*

$$\frac{\partial \kappa(G)}{\partial \ell(e)} = \frac{\partial}{\partial \ell(e)}(\kappa(G \setminus e) + \ell(e)\kappa(G/e)) = \frac{\partial}{\partial \ell(e)}\kappa(G \setminus e) + \frac{\partial}{\partial \ell(e)}(\ell(e)\kappa(G/e)) = \kappa(G/e).$$

□

We can also derive a differentiation rule for graphs with an available prime decomposition. Then, since prime decomposition is a partition of labels, we need only contract the edge in a single prime component.

**Identity 3.2.**

$$\frac{\partial \kappa(G)}{\partial \ell(e)} = \kappa(P_1/e)\prod_{i=2}^{n}\kappa(P_i),$$

where $P_1$ is the single prime component of $G$ containing $e$ and $P_i$ are the remaining $n-1$ prime components.

Along the same lines, we can derive a formula for the ratio of Kirchhoff polynomials composed of two graphs $G$ and $H$.

**Identity 3.3.** *Let the prime factorisation of the Kirchhoff polynomials $\kappa(G)$ and $\kappa(H)$ be $\kappa(G) = \prod_{i=1}^{n}\kappa(P_i)$ and $\kappa(H) = \prod_{j=1}^{m}\kappa(Q_j)$, then:*

$$\frac{\partial}{\partial \ell(e)}\frac{\kappa(G)}{\kappa(H)} = \frac{\prod_{i=2}^{n}\kappa(P_i)}{\kappa(Q_1)^2\prod_{j=2}^{m}\kappa(Q_j)}\left(\kappa(P_1/e)\kappa(Q_1) - \kappa(P_1)\kappa(Q_1/e)\right),$$

where $P_1$ and $Q_1$ are the prime components of $G$ and $H$, correspondingly, containing $e$.

*Derivation.*

$$\begin{aligned}\frac{\partial}{\partial e}\frac{\kappa(G)}{\kappa(H)} &= \frac{\kappa(P_1/e)\prod_{i=2}^{n}\kappa(P_i)\prod_{j=1}^{m}\kappa(Q_j) - \kappa(Q_1/e)\prod_{i=1}^{n}\kappa(P_i)\prod_{j=2}^{m}\kappa(Q_j)}{[\prod_{j=1}^{m}\kappa(Q_j)]^2}\\ &= \frac{\prod_{i=2}^{n}\kappa(P_i)\prod_{j=2}^{m}\kappa(Q_j)}{[\prod_{j=1}^{m}\kappa(Q_j)]^2}\left(\kappa(P_1/e)\kappa(Q_1) - \kappa(P_1)\kappa(Q_1/e)\right)\\ &= \frac{\prod_{i=2}^{n}\kappa(P_i)}{\kappa(Q_1)^2\prod_{j=2}^{m}\kappa(Q_j)}\left(\kappa(P_1/e)\kappa(Q_1) - \kappa(P_1)\kappa(Q_1/e)\right).\end{aligned}$$

□

It could be possible to further factorise the differentiated expressions since edge contraction could change graph connectivity.

Integrating a Kirchhoff polynomial with respect to a label $\ell(e)$ corresponding to an edge $e$ is equivalent to multiplication by $\ell(e)$ and edge relabelling in the corresponding graph as seen from Identity 3.4.

**Identity 3.4.**

$$\int \kappa(G)d\ell(e) = \ell(e)\kappa(G^{\ell(e)\leftarrow\frac{\ell(e)}{2}}) + C,$$

where $C$ is the integration constant and $\ell(e) \leftarrow \frac{\ell(e)}{2}$ denotes a relabelling operation replacing the label of $e$ by the same label divided by two, e.g. if the old label was $\ell(e) = r_1$ the new would be $\ell(e) = \frac{r_1}{2}$.

*Derivation.*

$$\begin{aligned}\int \kappa(G)d\ell(e) &= \int(\kappa(G \setminus e) + \ell(e)\kappa(G/e))d\ell(e) = \int \kappa(G \setminus e)d\ell(e) + \int \ell(e)\kappa(G/e)d\ell(e) =\\ &= \ell(e)\kappa(G \setminus e) + C_1 + \kappa(G/e)\int \ell(e)d\ell(e) = \ell(e)\kappa(G \setminus e) + C_1 + \frac{\ell(e)^2}{2}\kappa(G/e) + C_2 =\\ &= \ell(e)\left(\kappa(G \setminus e) + \frac{\ell(e)}{2}\kappa(G/e)\right) + C_1 + C_2 = \ell(e)\kappa(G^{\ell(e)\leftarrow\frac{\ell(e)}{2}}) + C.\end{aligned}$$

□

Note that integration does not change the factorisation properties of $G$ since its connectivity remains unchanged and that the labels remain unique unless $\frac{\ell(e)}{2}$ is already labelling another edge in the graph.

### 4 Generation of Kirchhoff polynomials

By GetPrimeDecomposition we denote the function taking a graph and returning its prime components, whose pseudocode can be found in (1). Also, we abbreviate directed graphs as digraphs.

**Algorithm 1** Recursive compressed generation of Kirchhoff polynomials through prime decomposition and edge deletion-contraction.

```
function C_R(G)
    if κ(G) == 0 then
        return 0
    end if
    if |V(G)| ≤ 2 then
        return GenKirchPolBaseCase(G)
    end if
    F ← []
    for all primeComponent ∈ GetPrimeDecomposition(G) do
        H ← GenKirchPolInPrimeComponent(primeComponent)
        F.append(H)
    end for
    return Multiply(F)                                    ▷ n-ary multiplication
end function

function GenKirchPolInPrimeComponent(G)
    if |V(G)| ≤ 2 then
        return GenKirchPolBaseCase(G)
    end if
    e ← GetEdgeForDelContr(G)
    kirchPolEdgeDelDigraph ← C_R(G \ e)
    kirchPolEdge ← GenKirchPolBaseCase(e)
    kirchPolEdgeContrDigraph ← C_R(G/e)
    return Add(kirchPolEdgeDelDigraph, Multiply(kirchPolEdge, kirchPolEdgeContrDigraph))
end function

function GenKirchPolBaseCase(G)
    if κ(G) == 0 then
        return 0
    end if
    if |V(G)| == 1 then
        return 1
    end if
    F ← []
    for all e ∈ E(G) do
        F.append(ℓ(e))
    end for
    return Add(F)                                         ▷ n-ary addition
end function
```

**Algorithm 2** Iterative compressed generation of Kirchhoff polynomials through prime decomposition, edge deletion-contraction, and change of variables.

```
function C_I(G)
    G.POINTER ← startPointer                                  ▷ every digraph has a unique pointer
    (P, R) ← GENIMPLICITKIRCHPOL(G)
    Q ← QUEUE()
    Q.ENQUEUE(startPointer)
    result ← []
    while Q not empty do
        currPointer ← Q.DEQUEUE()
        currExpr ← R.RETRIEVEEXPRESSION(currPointer)          ▷ currExpr is an expression tree
        result.APPEND((currPointer, ASSEMBLE(P, R, Q, currExpr)))   ▷ pass Q by reference
    end while
    return result
end function

function GENIMPLICITKIRCHPOL(G)
    Q ← QUEUE()
    Q.ENQUEUE(G)
    (P, R) ← ([], [])             ▷ P remembers already processed digraphs and R holds the results
    while Q not empty do
        H ← Q.DEQUEUE()
        primeComps ← GETPRIMEDECOMPOSITION(H)
        if H not prime OR κ(H) == 0 OR κ(H) == 1 then
            pointers ← GETPOINTERS(primeComps) ▷ list of primeComps's pointers; returns 0 or 1
if κ(H) == 0 or κ(H) == 1, respectively; calls base case
            expr ← MULTIPLY(pointers)
            R.APPEND((H.POINTER, expr))
        end if
        if κ(H) ≠ 0 AND κ(H) ≠ 1 then
            for all D ∈ primeComponents do
                if |V(D)| ≤ 2 then                                        ▷ base case
                    R.APPEND((D.POINTER, GENKIRCHPOLBASECASE(D)))
                else
                    if ∃ i: EQL(D,P_{i,0}) then         ▷ EQL() tests Kirchhoff polynomial equality
                        P_{i,1}.APPEND(D.POINTER)
                    else                                           ▷ digraph not investigated yet
                        P.APPEND((D, []))
                        e ← GETEDGEFORDELCONTR(D)
                        (DDelE, E, DContrE) ← (D \ e, GENKIRCHPOLBASECASE(e), D/e)
                        expr ← ADD(DDelE.POINTER, MULTIPLY(E, DContrE.POINTER))
                        R.APPEND((D.POINTER, expr))
                        Q.ENQUEUE(DDelE, DContrE)
                    end if
                end if
            end for
        end if
    end while
    P ← REMOVEUNMATCHEDDIGARPHS(P)
    return (P, R)
end function
```

As in $C_R$, given a graph $G$, $C_I$ alternates between prime decomposition and edge deletion-contraction to reduce the Kirchhoff polynomial generation problem to several smaller ones. However, in $C_I$ a unique pointer is associated to every graph under study, and $C_I$ adds the reduced graphs to a

**Algorithm 2** Continued.

```
function ASSEMBLE(P,R, Q, currExpr)
    if currExpr is MULTIPLY or ADD then
        newChildren ← []
        for all childExpr ∈ currExpr.CHILDREN do
            newChildren.APPEND(ASSEMBLE(P, R, Q, childExpr))
        end for
        currExpr.CHILDREN ← newChildren
    else if currExpr is a POINTER then
        if ∃i : currExpr == P_{i,0}.POINTER then
            Q.ENQUEUE(currExpr)
        else if ∃p and j : p ∈ P_{j,1} and currExpr == p then
            currExpr ← P_{j,0}.POINTER
        else
            newExpr ← R.RETRIEVEEXPRESSION(currExpr)
            return ASSEMBLE(P,R, Q, newExpr)
        end if
    end if
    return currExpr
end function
```

queue for further reduction, simultaneously remembering the partial expression tree they participate in. A *partial expression tree* is such a tree in which leaves could correspond to Kirchhoff polynomials. For example, the deletion-contraction identity from Equation 6 provides a partial expression tree in which two leaves are Kirchhoff polynomials, $\kappa(G \setminus e)$ and $\kappa(G/e)$, and one is an edge label $\ell(e)$. Thus this tree can be remembered after substituting $\kappa(G \setminus e)$ and $\kappa(G/e)$ with their pointers, and the graphs corresponding to these Kirchhoff polynomials can be taken for further reduction. Additionally, during the reduction procedure every prime component is compared to all previously encountered prime components using the criteria from Corollary 3.2 and Corollary 3.7. If the equality of a prime component's Kirchhoff polynomial to a previously considered graph's Kirchhoff polynomial cannot be certified, then the prime component is taken for further reduction. In contrast, if a prime component and a previously encountered graph $H$ have equal Kirchhoff polynomials, then the reduction of the prime component is discontinued and its Kirchhoff polynomial is substituted with the pointer of $H$, thus marking the identity. Algebraically this is equivalent to a change of variables – substituting identical parts of the Kirchhoff polynomial with identical symbols and explicitly generating them only once (see Figure 2d for an example). The reduction procedure again continues until graphs are reduced to a single vertex or a single edge and produces a set of partial expression trees.

The partial expression trees are then assembled. The assembly starts from the given graph $G$ (with pointer $S$) and its partial expression tree which is sequentially merged with the partial expression trees of its reduced graphs. The merging proceeds if the current reduced graph has not been matched with another graph with identical Kirchhoff polynomial. If a match is present, then the partial expression tree of the reduced graph is substituted with the pointer, e.g. $X$, (as a variable) of a predetermined graph with an equal Kirchhoff polynomial encountered during the reduction procedure (could be the current graph itself) and merging is discontinued. Simultaneously, another assembly is initiated starting from $X$ and its corresponding graph to obtain a forest of expressions marked with the pointers of the initializing graphs as in Figure 2d. This forest of expression trees corresponds to a set of Kirchhoff polynomials, which after being substituted into each other give rise to the complete Kirchhoff polynomial of the given graph $G$.

Note that substitution is unnecessary when evaluating the Kirchhoff polynomial for a given set of edge label values. There exists a sequence obtainable in linear time in which the expression trees from the forest can be evaluated such that there are no uncalculated pointer variables during the evaluation. The reason is that the expression trees in the forest can be thought of as arranged in a directed acyclic graph, with vertices being the trees themselves and edges being the change of variables directed relations, which can always be topologically sorted.

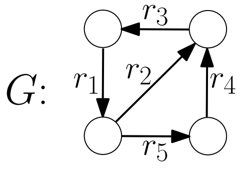


Figure S1: Example graph $G$ used in Figure 2.

(a)

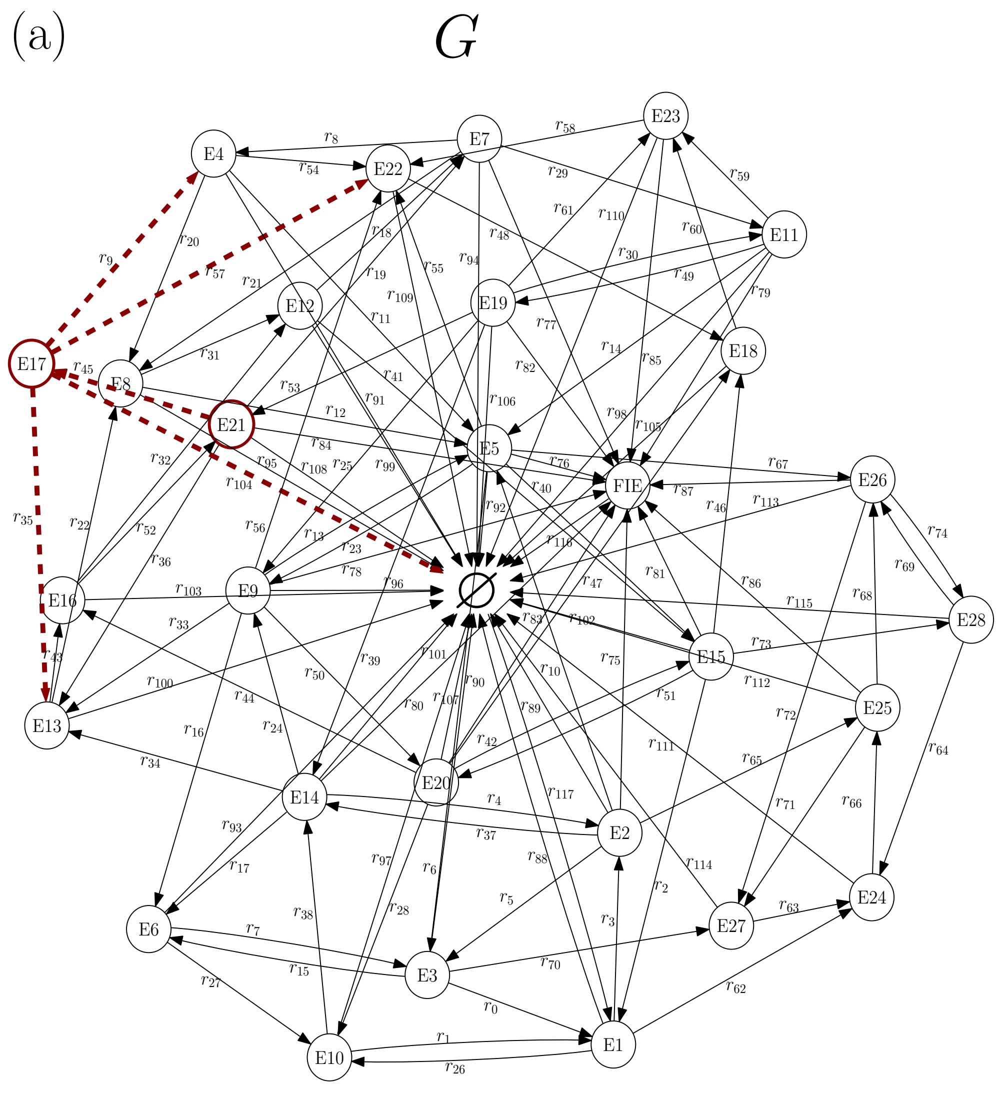


(b)

$$\frac{x^{SS}_{E21}}{x^{SS}_{E17}} = \frac{\frac{\kappa_{v_{E21}}(G)}{\cancel{\kappa_{v_{\varnothing}}(G)}}}{\frac{\kappa_{v_{E17}}(G)}{\cancel{\kappa_{v_{\varnothing}}(G)}}} = \frac{\kappa_{v_{E21}}(G)}{\kappa_{v_{E17}}(G)} = \frac{r_9+r_{35}+r_{104}+r_{57}}{r_{45}}$$

Figure S2: Simplification of expressions of Kirchhoff polynomials in the coarse-grained representation. (a) A graph $G$ for the PHGS catalytic cycle (see COX in Table S1) from (3) and (b) the steady-state ratio of species $E21$ and $E17$ obtained through simplification of the coarse-grained representation. Coloured vertices (arrows) denote species of interest (reactions contained in the simplified ratio).

#### 5 Description of used heuristics

Box 1 Heuristics.

**i.) Edges $E'(G)$:**

0. A single randomly selected edge $e$, $E'(G) = \{e\}$.
1. All edges, $E'(G) = E(G)$.
2. Edges participating in the longest simple cycle.
3. The $|E(G)|/n, n = 3$ edges participating in the largest number of simple cycles.

**ii.) Branch:**

0. Edge deleted graph, $G \setminus e$.
1. Edge contracted graph, $G/e$.
2. Edge deleted and edge contracted graphs, $(G \setminus e, G/e)$.

**iii.) Components:**

0. Strongly connected components.
1. Prime components.

**iv.) Optimality criterion:**

0. Largest number of components.
1. Largest component (in terms of number of vertices) is smallest.
2. Largest component (in terms of number of edges) is smallest.
3. Smallest total complexity (number of arborescences) of the components.
4. Smallest total complexity with the largest (same) number of components.
5. Largest number of components with the (same) smallest total complexity.

We consider each heuristic to be composed of a four-step procedure. We propose several choices of *sub-heuristics* for each step. The set of all heuristics of interest contains all combinations of these sub-heuristics at the different steps of the procedure. Formally, we use the same heuristic $\mathcal{H}$ for the whole reduction process of a given input graph and we explore all 108 combinations of sub-heuristics, which we call *connectivity-informed heuristics*. In addition, one heuristic, denoted by $\mathcal{H} = 0 * **$, selects a random edge, uninformed by the graph connectivity.

More precisely, the procedure is (see also Box 1 for a more concise description):

(i) Pre-select a subset of edges $E'(G) \subseteq E(G)$.

This subset could be a single randomly picked edge (leading to a random *connectivity-uninformed heuristic*), all edges, or a subset of edges connected to the graph cycle structure (aiming to break open many or large cycles).

(ii) Apply edge deletion-contraction to $G$ for each $e \in E'(G)$ and decide which branch of the deletion-contraction tree to consider, i.e. the edge deleted graph $G \setminus e$, the edge contracted graph $G/e$, or both $(G \setminus e, G/e)$.

This is required since edge contraction could also lead to edge deletions and thus to further prime decomposition.

(iii) Choose whether to decompose the graphs in the considered edge deletion-contraction branch(es) to strongly connected components or to prime components.

SCC decomposition alone leads to Kirchhoff polynomial factorisation which is not guaranteed to be prime. Yet we include it as a sub-heuristic due to recent results in strong connectivity allowing to retrieve all strong bridges (4), the total number of SCCs, and the size of the largest and of the smallest SCCs obtained after edge deletion in linear time (5). Note that, in order to have comparable running times when decomposing into prime components and SCCs, we naively delete-contract each considered edge and do not employ the mentioned recent advancements.

(iv) Calculate a score on the graph decomposition and pick the edge producing an optimal score.

The score is based on the number, size distribution (in terms of number of vertices or edges), and *complexity* (number of spanning trees) of components in the selected branch(es) of the deletion-contraction tree. The scores from the decompositions of $G \setminus e$ and $G/e$ are summed when both branches are taken into account. We choose the edge whose deletion leads to a decomposition in which there are the largest number of SCCs/prime components, the largest component is the smallest, the total complexity is the smallest, or there are the largest number of components with the smallest total complexity.

We describe each heuristic by four integers, where each integer marks the choice of a sub-heuristic. See Box 1 for the identifiers of each sub-heuristic we consider. For example, the heuristic $\mathcal{H} = 2205$ translates to a procedure in which we:

(i) **2:** Find the longest simple cycle $S$ in the input graph $G$ and take its edges $E(S)$.

(ii) **2:** For every $e \in E(S)$ we apply deletion-contraction to obtain the graphs from both branches of the deletion-contraction tree, $G \setminus e$ and $G/e$.

(iii) **0:** Obtain the strongly connected components for every $G \setminus e$ and $G/e$ and add them to a list $p_e$.

(iv) **5:** Pick the edge producing the list $p_e$ with the largest length and return the edge $e$. If there are several lists having the same length, we pick the one with the smallest total sum of component complexities and return the corresponding edge $e$.

### 6 Collection of graphs

Table S1: A collection of example graphs, ordered by their complexity (number of spanning trees). Shown are the graph aliases (under $G$), number of vertices $|V|$, number of edges $|E|$, number of spanning trees $|\mathsf{spt}(G)|$, and a short description of the model from which the graph was extracted. Note that during graph extraction all algebraic expressions in edge labels are taken as uninterpreted symbols. Additionally, some models may not satisfy all linear framework requirements, e.g. the time-scale separation assumption, or assume that equilibrium steady states are biologically relevant due to lack of evidence for energy dissipation or simpler algebraic derivations. Still, endowing their graphs with non-equilibrium Laplacian dynamics and deriving their Kirchhoff polynomials, could facilitate the understanding of their steady-state information processing capabilities and aims to inspire future applications employing the theory and algorithms developed in this work.

| $G$ | $\|V\|$ | $\|E\|$ | $\|\mathsf{spt}(G)\|$ | Description |
|---|---|---|---|---|
| COLE1 | 6 | 10 | 26 | Kinetic scheme of the ColE1 plasmids non-equilibrium replication control mechanism from (6). |
| AMPAR | 7 | 14 | 30 | State transition diagram of the AMPA receptor trafficking model from (7). This model is part of a signalling pathway model of corticostriatal spines that express D1-type dopamine receptors. |

Table S1 Continued:

| $G$ | $\lvert V\rvert$ | $\lvert E\rvert$ | $\lvert\mathsf{spt}(G)\rvert$ | Description |
|---|---|---|---|---|
| MDH | 9 | 18 | 141 | Proposed non-equilibrium kinetic mechanism for the reaction cycle of *M.methylotrophus* methanol dehydrogenase (MDH) with ammonium as activator from (8). |
| ACTMYO | 10 | 20 | 356 | State transition diagram detailing the interaction of actin and myosin from (9). The diagram is part of a model including calcium binding to troponin C and two configurations of tropomyosin. |
| KNF33 | 9 | 24 | $1,728$ | General allosteric model of ATP hydrolysis and competitive inhibition at two binding sites (10). Allostery is usually assumed to be happening at thermodynamic equilibrium, partly because of the simpler steady state derivations (11). However, non-equilibrium allosteric models have also been of interest (12). |
| SHPIL | 10 | 26 | $4,560$ | Model describing early IL-6 induced signaling. Model M0 from (13) in which time-scale separation is not assumed. |
| GR | 13 | 32 | $5,057$ | Scheme for the catalytic mechanism of glutathione reductase (GR) from (14). Rapid equilibrium was assumed in the original model to simplify kinetic flux derivations. |
| PHO5 | 12 | 35 | $53,376$ | A non-equilibrium linear framework model for the regulation of yeast PHO5 gene from (15). |
| RND | 14 | 36 | $69,120$ | Random Ter-Ter mechanism from (16). In the original model, rapid equilibrium was assumed for some reactions. |
| TF | 25 | 49 | $1,549,872$ | Largest strongly connected component of the transcription factor network of *Saccharomyces cerevisiae* extracted from (17). The network has not originally been endowed with Laplacian dynamics but deriving its Kirchhoff polynomial could be instructive when analysing its topological properties, e.g. when calcualating the spanning tree centrality of its vertices (18). |
| HC4 | 16 | 60 | $42,467,328$ | Four dimensional hypercube graph rooted at a vertex (the specific rooting does not matter due to symmetry). It represents a generic linear framework transcription regulation model for a gene with a promoter containing four transcription factor binding sites (19). |
| COXD | 30 | 117 | $12,254,915,821,568,674$ | Example COX rooted at the environment vertex, i.e. $\mathsf{rt}_{\varnothing}(\text{COX})$, which is the graph in the denominator of the steady-state expression for COX. |

Table S1 Continued:

| $G$ | $\lvert V\rvert$ | $\lvert E\rvert$ | $\lvert\mathsf{spt}(G)\rvert$ | Description |
|---|---|---|---|---|
| COX | 30 | 118 | $24,509,831,643,137,316$ | Scheme of the catalytic cycle of PGHS considering inhibition by NSAID from (3), Figure 1 (model obtained by personal communication with Alexey Goltsov, November, 2015). |

### 7 Kirchhoff polynomial generation results

We obtained the compression and average running time (from 10 runs) for each example graph from Table S1 using the 108 connectivity-informed heuristics with each algorithm, $C_R$ and $C_I$. The connectivity-uninformed random heuristic $\mathcal{H} = 0 * **$ was run 20 times on each low complexity example for both $C_R$ and $C_I$.

To understand the heuristics' influence, we contrasted the algorithms' performance using the random uninformed heuristic $\mathcal{H} = 0 * **$ with the informed heuristics incorporating directed graph connectivity information. Performance results are shown in Figure S5. They indicate that several runs with the uninformed heuristic suffice to generate highly compressed Kirchhoff polynomials in short time for examples of low complexity, possibly due to the lack of computational overhead required for the informed heuristics. However, for more complex graphs, the random heuristic quickly deteriorates, becoming orders of magnitude worse than the informed heuristics. This behaviour is expected since compressibility depends on graph connectivity and it becomes less probable to randomly pick an appropriate set of edges for deletion-contraction in larger graphs without any connectivity information. Additionally, note that running time and compression correlate inversely and that the correlation becomes more pronounced when the model complexity increases.

To assess the relative efficiency of the heuristics applied on the collection of less complex examples we normalized the performance measures over all heuristics separately for each example and divided them into groups. The resulting box plots for algorithm $C_I$ can be seen in Figure S3. The results for $C_R$ are similar and are presented in Figure S4. We observe that some sub-heuristics, on average, perform better both in running time and compression on the collection of examples. We performed the non-parametric Kruskal-Wallis H-test which showed that in every group there was at least one choice for a sub-heuristic dominating the others, with the exception of the compression results of group "(iii) Components" in which no sub-heuristic dominates. Post-hoc comparisons of sub-heuristic choices using the Wilcoxon signed-rank test revealed that focusing deletion-contraction to edges relevant to the cycle structure of the graphs (sub-heuristics (i) **2** and (i) **3**), considering the edge deleted graphs (sub-heuristic (ii) **0**) and strongly connected components (sub-heuristic (iii) **0**), and not picking sub-heuristic (iv) **2** leads to significantly shorter running time and larger compression on average (see Table S2 and Table S3 for results). A limitation in the significance analysis is the different skewness of the data and the dependency between the sub-heuristics, e.g. a good optimality criterion choice cannot remedy a bad edge subset choice. Thus we cannot conclude that a given heuristic, e.g. $\mathcal{H} = 3002$, always leads to fast running time and good compression.

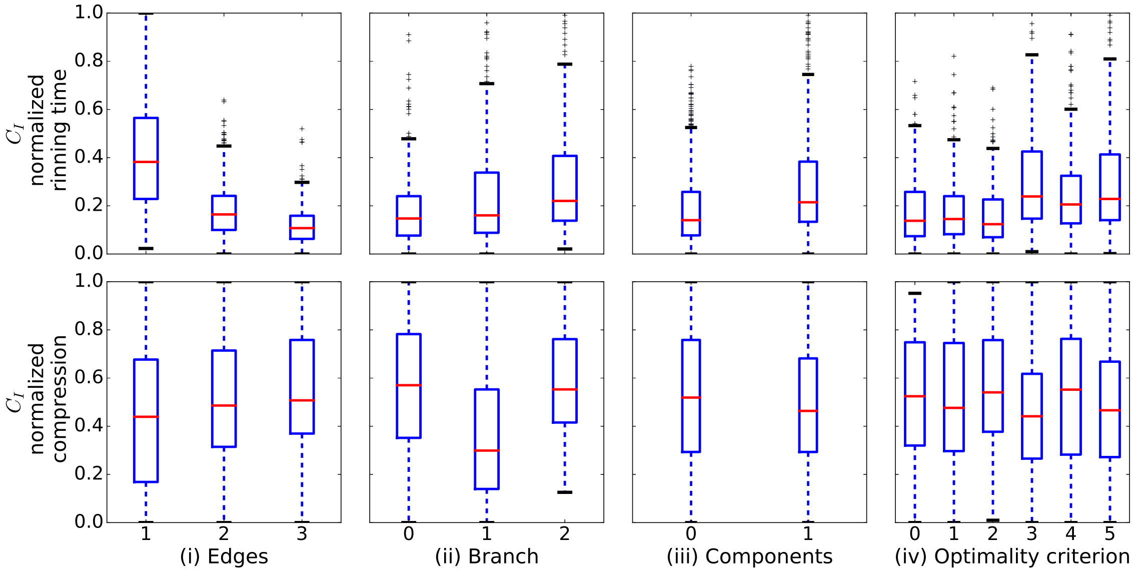


Figure S3: Comparison of the normalised running time and compression distributions on the set of less complex examples for algorithm $C_I$ grouped by the different choices of each sub-heuristic - edges, branch, components, and optimality criterion. The running time and compression were normalized from 0 to 1, using the formula $\frac{x-min}{max-min}$, where $x$ is the running time/compression for every heuristic and example, $min$ is the shortest running time/smallest compression and $max$ is the longest running time/largest compression among all heuristics for the particular example.

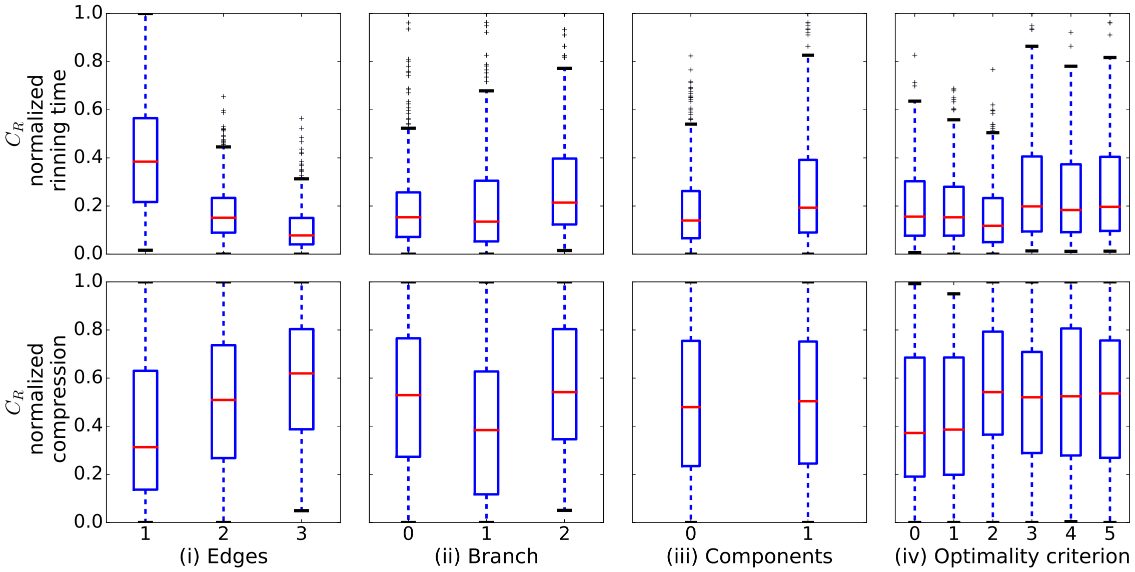


Figure S4: Comparison of the normalised running time and compression distributions on the set of less complex examples for algorithm $C_R$ grouped by the different choices of each sub-heuristic - edges, branch, components, and optimality criterion.

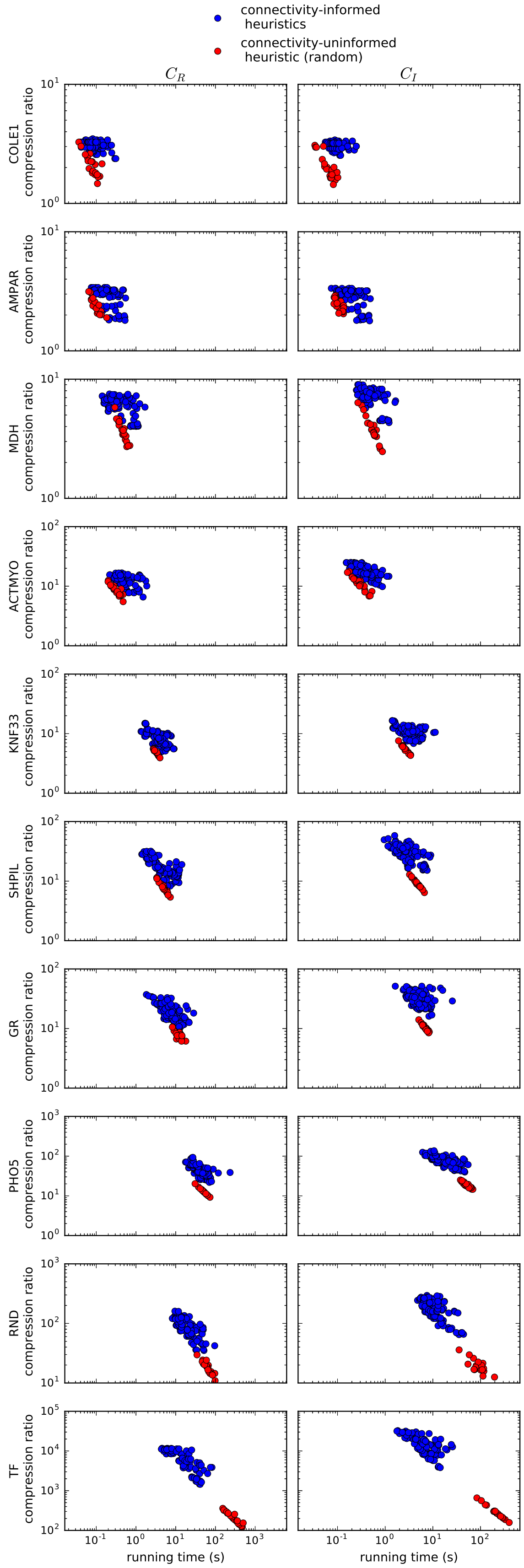


Figure S5: Scatter plots of running time versus compression obtained with each of the two algorithms $C_R$ (left column) and $C_I$ (right column) on the collection of less complex examples. Each row corresponds to a graph example and the examples are sorted by increasing graph complexity. Blue points represent performance results for each of the 108 connectivity-informed heuristics and red points mark the results for the 20 runs of the uninformed random heuristic $\mathcal{H} = 0 * **$.

Table S2: Significance levels (p-values) of the different choices of sub-heuristics for the running time results of algorithm $C_I$ on the less complex set of graph examples. The non-parametric Kruskal-Wallis H-test was applied along with post hoc pair comparisons using the Wilcoxon signed-rank test to determine whether the samples originate from the same distribution.

| Test | (i) Edges | | (ii) Branch | | (iii) Components | | (iv) Optimality | |
|---|---|---|---|---|---|---|---|---|
| | pair | p-value | pair | p-value | pair | p-value | pair | p-value |
| Kruskal-Wallis H-test | – | 1.37e-88 | – | 2.91e-14 | – | 1.30e-16 | – | 1.14e-19 |
| Wilcoxon signed-rank test | (1, 2) | 3.87e-55 | (0, 1) | 5.39e-07 | (0, 1) | 5.50e-72 | (0, 1) | 7.35e-01 |
| | (1, 3) | 4.10e-58 | (0, 2) | 1.64e-34 | | | (0, 2) | 4.94e-03 |
| | (2, 3) | 3.38e-28 | (1, 2) | 4.08e-09 | | | (0, 3) | 8.47e-21 |
| | | | | | | | (0, 4) | 4.99e-17 |
| | | | | | | | (0, 5) | 5.30e-20 |
| | | | | | | | (1, 2) | 2.60e-03 |
| | | | | | | | (1, 3) | 6.47e-20 |
| | | | | | | | (1, 4) | 1.19e-14 |
| | | | | | | | (1, 5) | 3.59e-19 |
| | | | | | | | (2, 3) | 3.59e-28 |
| | | | | | | | (2, 4) | 3.39e-23 |
| | | | | | | | (2, 5) | 1.26e-27 |
| | | | | | | | (3, 4) | 4.61e-10 |
| | | | | | | | (3, 5) | 1.44e-02 |
| | | | | | | | (4, 5) | 5.38e-07 |

Table S3: Significance levels (p-values) of the different choices of sub-heuristics for the compression results of algorithm $C_I$ on the less complex set of graph examples.

| Test | (i) Edges | | (ii) Branch | | (iii) Components | | (iv) Optimality | |
|---|---|---|---|---|---|---|---|---|
| | pair | p-value | pair | p-value | pair | p-value | pair | p-value |
| Kruskal-Wallis H-test | – | 4.15e-05 | – | 4.55e-33 | – | 1.01e-01 | – | 2.13e-02 |
| Wilcoxon signed-rank test | (1, 2) | 9.08e-07 | (0, 1) | 9.41e-18 | (0, 1) | 1.78e-04 | (0, 1) | 7.75e-01 |
| | (1, 3) | 4.47e-08 | (0, 2) | 9.03e-01 | | | (0, 2) | 4.87e-01 |
| | (2, 3) | 5.14e-02 | (1, 2) | 8.67e-36 | | | (0, 3) | 1.77e-03 |
| | | | | | | | (0, 4) | 4.61e-01 |
| | | | | | | | (0, 5) | 7.73e-02 |
| | | | | | | | (1, 2) | 2.85e-02 |
| | | | | | | | (1, 3) | 8.33e-03 |
| | | | | | | | (1, 4) | 5.49e-01 |
| | | | | | | | (1, 5) | 2.01e-01 |
| | | | | | | | (2, 3) | 2.35e-10 |
| | | | | | | | (2, 4) | 7.08e-01 |
| | | | | | | | (2, 5) | 5.21e-05 |
| | | | | | | | (3, 4) | 7.52e-08 |
| | | | | | | | (3, 5) | 8.66e-04 |
| | | | | | | | (4, 5) | 8.69e-05 |

### 8 Calculation of the position-steepness regions

To obtain boundaries of position-steepness regions we strictly follow the definitions and methods described in (19), with the exception of i) gene regulatory function (GRF) derivation, for which we apply the methods and algorithms developed in this work and ii) the boundary expansion algorithm, to which we apply changes that we empirically found to be more effective for the expansion in models with large parameter spaces.

#### 8.1 Compressed GRF derivation

We derive the symbolic non-equilibrium GRFs for hypercube linear framework models with $n = 2, 3, 4$ binding sites using the compressed Kirchhoff polynomial generation algorithm $C_I$ with heuristic 1001 implemented in *KirchPy*. We exemplify the compressed derivation results for the $n = 3$ sites model, which can be seen in Figure S6, since the symbolic GRF of the $n = 4$ sites model is too lengthy to include despite its 2000 times compression.

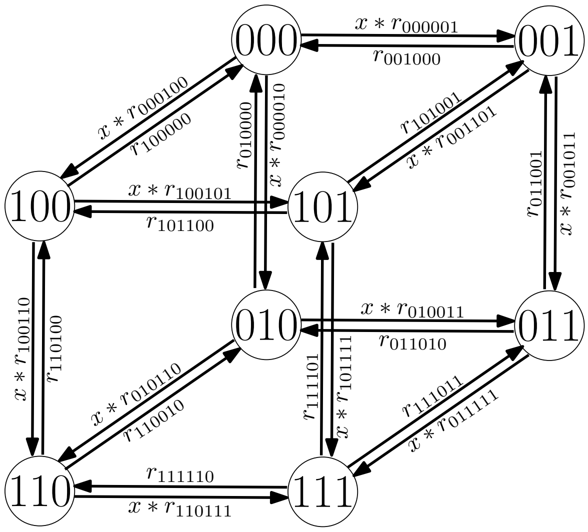


Figure S6: A linear framework model for a transcription factor (TF) binding to $n = 3$ DNA sites. The corresponding cube graph contains 8 vertices denoting DNA microstates, where each microstate is a pattern of bound TFs. Patterns are encoded in the vertex labels. For example, vertex label 101 denotes a DNA microstate, in which the first and last binding sites are bound to a TF, while the site between them is free. Edges denote transitions between microstates. Edge labels contain transition rates with $x$ standing for TF concentration and $r$s are rate constants; $r$'s subscripts name the corresponding transitions, e.g. $r_{101001}$ is in the label of the transition edge between microstates 101 and 001.

Considering graph $G$ from Figure S6 and the all-or-nothing expression strategy, we express the steady-state probability of the all-bound microstate 111, denoted as $p_{111}$, as a function of TF concentration $x$:

$$p_{111}(x) = \frac{\kappa_{v_{111}}(G)}{\sum_{v_\xi \in V(G)} \kappa_{v_\xi}(G)},$$

where $\xi$ marks TF binding patterns.

Observe that all rooted polynomials of $G$, $\kappa_{v_\xi}(G)$, are isomorphic to each other. Thus we can obtain mappings between the labels of, e.g. $\kappa_{v_{111}}(G)$, and the labels of any other $\kappa_{v_\xi}(G)$. We define a *template* function `TemplateHypercube3` taking a set of labels and returning the Kirchhoff polynomial $\kappa_{v_{111}}(G)$ with labels substituted according to the isomorphic mapping. In consequence, due to the symmetry of hypercube graphs, we need only generate a template Kirchhoff polynomial from a single rooted graph and call it with the appropriate labels, in order to evaluate any rooted polynomial. The resulting code used for the derivation of GRFs is shown in Listing S1. It produces symbolic GRFs in terms of the TF concentration variable such as those in Listing S2. In contrast to the numerical approach taken in (19), we extract the position and steepness features from these symbolic GRFs by taking symbolic derivatives, which is more efficient and less prone to numerical errors.

Note that identifying symmetries and isomorphism in LFMs can reduce Kirchhoff polynomial generation time and yield a shorter representation. However, unlike our compression methods, isomorphism does not offer speed-ups to numerical evaluation since it does not reduce the number of arithmetic operations.

Listing S1: A *Mathematica* function for the derivation of GRFs with n=3 binding sites and the all-or-nothing expression strategy as a function of TF concentration $x$.

```
AllOrNothing3[parVect_]:=Module[{r000001,r000010,r000100,r001000,r001011,r001101,
    r010000,r010011,r010110,r011001,r011010,r011111,r100000,r100101,r100110,r101001,
    r101100,r101111,r110010,r110100,r110111,r111011,r111101,r111110,root010,root011,
    root001,root000,root111,root110,root100,root101,response},

(* r -- transition rate between two states,e.g. r001101 denotes the label associated
    to edge 001->101 *)

(* non-deminsionalisation of rate constants *)
r100000 = 1;
r000100 = 1;

(* 22 transition rates as free variables *)
r000001 = parVect[[1]];
r000010 = parVect[[2]];
r001000 = parVect[[3]];
r001011 = parVect[[4]];
r001101 = parVect[[5]];
r010000 = parVect[[6]];
r010011 = parVect[[7]];
r010110 = parVect[[8]];
r011001 = parVect[[9]];
r011010 = parVect[[10]];
r011111 = parVect[[11]];
r100101 = parVect[[12]];
r100110 = parVect[[13]];
r101001 = parVect[[14]];
r101100 = parVect[[15]];
r101111 = parVect[[16]];
r110010 = parVect[[17]];
r110100 = parVect[[18]];
r110111 = parVect[[19]];
r111011 = parVect[[20]];
r111101 = parVect[[21]];
r111110 = parVect[[22]];

(* x -- transcription factor concentration taken as a symbolic variable *)
(* root -- Kirchhoff polynomials rooted at each of the vertices of the cube model,
    obtained by calling the template function with the appropriate input determined by
     the isomorphism between rooted graphs in G *)

root010 = TemplateHypercube3[r110100,r110111*x,r001101*x,r001000,r000100*x,r000010*x,
    r000001*x,r101100,r101111*x,r101001,r111110,r111101,r110010,r111011,r011111*x,
    r011001,r011010,r100110*x,r100101*x,r100000,r001011*x];
root011 = TemplateHypercube3[r001000,r001101*x,r110100,r110010,r010000,r010011*x,
    r010110*x,r100000,r100101*x,r100110*x,r101001,r101100,r001011*x,r101111*x,r111101,
    r111110,r111011,r000001*x,r000100*x,r000010*x,r110111*x];
root001 = TemplateHypercube3[r000100*x,r000010*x,r111110,r111101,r101100,r101001,
    r101111*x,r110100,r110010,r110111*x,r010000,r010110*x,r000001*x,r010011*x,r011010,
    r011111*x,r011001,r100000,r100110*x,r100101*x,r111011];
root000 = TemplateHypercube3[r010110*x,r010011*x,r101111*x,r101100,r100110*x,r100000,
    r100101*x,r111110,r111011,r111101,r011010,r011111*x,r010000,r011001,r001011*x,
    r001101*x,r001000,r110010,r110111*x,r110100,r101001];
root111 = TemplateHypercube3[r101100,r101001,r010000,r010110*x,r110100,r110111*x,
    r110010,r000100*x,r000001*x,r000010*x,r001101*x,r001000,r101111*x,r001011*x,
    r011001,r011010,r011111*x,r100101*x,r100000,r100110*x,r010011*x];
root110 = TemplateHypercube3[r111101,r111011,r000001*x,r000100*x,r100101*x,r100110*x,
    r100000,r001101*x,r001011*x,r001000,r011111*x,r011001,r111110,r011010,r010011*x,
    r010000,r010110*x,r101111*x,r101001,r101100,r000010*x];
root100 = TemplateHypercube3[r101111*x,r101001,r010011*x,r010110*x,r110111*x,r110100,
    r110010,r011111*x,r011001,r011010,r001101*x,r001011*x,r101100,r001000,r000001*x,
    r000010*x,r000100*x,r111101,r111011,r111110,r010000];
root101 = TemplateHypercube3[r100110*x,r100000,r011010,r011111*x,r111110,r111101,
    r111011,r010110*x,r010000,r010011*x,r000100*x,r000010*x,r100101*x,r000001*x,
```

```
    r001000,r001011*x,r001101*x,r110100,r110010,r110111*x,r011001];

response = (root111)/Collect[root010+root011+root001+root000+root111+root110+root100+
    root101,x];

Return[response];
];

TemplateHypercube3[G0_,G1_,G10_,G11_,G12_,G13_,G14_,G15_,G16_,G17_,G18_,G19_,G2_,G20_
    ,G3_,G4_,G5_,G6_,G7_,G8_,G9_]:=Module[{psi61,psi34,psi75,start},
(* template expression -- Kirchhoff polynomial rooted at a vertex *)

psi61 = Collect[((G8+G7+G6)*(G1+G2)+G0*(G8+G7)),x];
psi34 = Collect[(psi61*(G14+G13)+G12*((G7+G6)*(G1+G2)+G0*G7)),x];
psi75 = Collect[((G20+G18)*(G0+G2)+G1*G20),x];
start = Collect[(G10*((G3+G4+G5)*((G12+G14+G13)*(((G15+G16)*G6+G7*G16)*G18+G19*G15*G6
    )*G2+G8*(((G19+G18)*(G0+G2)+G1*G19)*G15*G13+G16*G18*((G13+G14)*G2+G0*G13)))+G20*(
    psi34*G16*G5+G15*((G1+G0+G2)*(G4+G5)*G8*G13+G6*(G13+G12+G14)*((G5+G4)*G2+G1*G5))))
    +G19*((G4+G3+G5)*G11*((psi61*G17+G15*((G8+G6)*G2+(G0+G1)*G8))*G13+G12*(G15+G17)*G6
    *G2)+G9*(psi34*G17*G5+G15*((G14+G12+G13)*(G3+G5)*G6*G2+G8*(G0+G1+G2)*((G3+G5)*G13+
    G14*G5))))+(G16+G15+G17)*(G4*G11*(psi75*G8*G13+G6*((G20+G18)*(G13+G12)*G2+G1*G20*
    G13))+((G9+G11)*((G6+G8)*G13+G12*G6)+G14*(G6+G8)*G9)*(((G20+G18)*G2+G1*G20)*G5+G3*
    G18*G2)+G0*G8*((G9+G11)*((G20+G18)*G5+G3*G18)*G13+G14*(G20+G18)*G9*G5))+G7*((G14+
    G12+G13)*G9*((G1+G0+G2)*(G16+G17)*G20*G5+G18*((G17+G16)*(G5+G3)*G2+G0*G17*G5))+G11
    *((psi75*(G4+G5)+G3*G18*(G0+G2))*G17*G13+G16*((G3+G4+G5)*(G12+G13)*G18*G2+G20*(G1+
    G0+G2)*((G4+G5)*G13+G12*G5))))),x];

Return[start];
];
```

#### 8.2 Boundary expansion algorithm

In (19) the boundaries of position-steepness regions are expanded using a biased sampling algorithm (BSA). BSA produces an initial region by randomly sampling GRF parameters from a specified parameter box, calculating the position and steepness features of the resulting GRFs, and computing the enclosing boundary. The initial boundary is successively expanded by randomly changing the parameters of GRFs on the current boundary until the area of the region does not increase any further.

We observed that BSA frequently gets stuck in local minima and does not have a satisfactory performance to expand the boundary for the $n = 4$ model with 62 free variables. Hence, in an attempt to speed-up the convergence of boundary expansion, we alternate between BSA and a simple optimisation procedure. Namely, we i) take a GRF with position-steepness coordinates at the boundary (or close to the boundary, but inside the region), ii) pick a position-steepness point outside the boundary in the direction of desired expansion, and iii) minimize the Euclidean distance between the points in terms of the GRF's parameters using *Matlab*'s function `fmincon` subject to the parameter box constraints. Additionally, we modify the step in BSA in which GRF parameters are changed by allowing a subset of the parameters, e.g. $10 - 30$, to be "mutated" at a time. This adjustment is needed to obtain mutated GRFs in the position-steepness neighbourhood of the original GRF because big changes in many parameters can significantly alter GRF shape, which we observed to overwhelmingly produce GRFs inside the boundary.

#### 8.3 Results

As a result of the compressed GRF generation and the changes to the boundary expansion algorithm we obtained the boundaries in Figure S7. The symbolic form of specific GRFs lying on the boundaries can be found in Listing S2.

Listing S2: Unnormalised GRFs corresponding to points on the boundary of Figure S7.

```
%symbolic GRF for Figure S7, a):
GRF_a(x) = (3.111898366*x^2 + 1088.2653466091372*x^3)/(0.9798786583499999 +
    3.0441335554862703*x + 4.660564433985895*x^2 + 1088.2653466091372*x^3)

%symbolic GRF for Figure S7, b):
GRF_b(x) = (4.506992106647366*10^10*x^3 + 8.838579100646242*10^12*x^4 +
    3.494151550118564*10^14*x^5 + 2.006924322139229*10^16*x^6 +
    6.107156670959329*10^17*x^7)/(3.938715616171477*10^9 + 1.0710384966602333*10^10*x
```

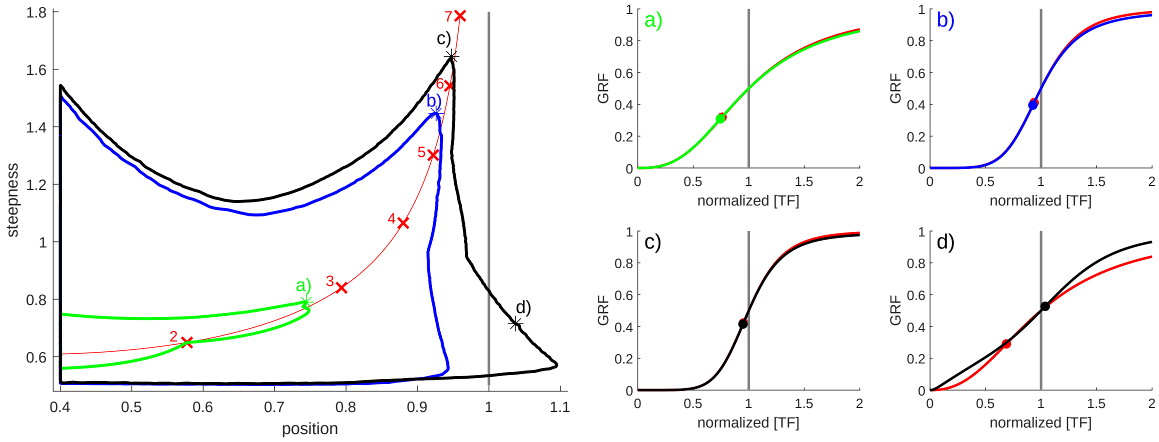


Figure S7: **(left)** Position-steepness regions for non-equilibrium gene response functions corresponding to models with $n = 2$ (green), $n = 3$ (blue), $n = 4$ (black) transcription factor binding sites. The boundaries are obtained for the all-or-nothing expression strategy by sampling parameter values in the interval $[10^3, 10^{-3}]$. The four points on the boundaries — a), b), c), and d)— correspond to the GRFs on the right-hand-side of the figure. The *Hill line* defined by the position-steepness loci of Hill functions with coefficients ranging from 1.5 to 7 is shown in red; loci corresponding to integer Hill coefficients are marked with red crosses and numbers. The position asymptote of the Hill line is marked in grey. **(right)** GRF value (probability of the 111 microstate) as a function of normalised TF concentration for the four points lying on the boundaries in the left-hand-side of the figure. The correspondence between position-steepness points and GRFs is marked by colour and letters. Red curves are Hill functions with identical steepness. Dots denote the position of maximal steepness. Note that TF concentration is normalised such that the grey line at 1 crosses the GRFs at their half-maximal value.

```
    + 2.443283916713498*10^11*x^2 + 1.5688345854462803*10^12*x^3 +
    1.762963633037459*10^13*x^4 + 4.326523028160712*10^14*x^5 +
    2.108616283332164*10^16*x^6 + 6.107156670959329*10^17*x^7)

%symbolic GRF for Figure S7, c):
GRF_c(x) = (1.2224524729375644*10^39*x^4 + 5.319185045157951*10^40*x^5 +
    6.489224875394841*10^41*x^6 + 4.0133242397037175*10^42*x^7 +
    1.8140472985591478*10^43*x^8 + 6.423734524550449*10^43*x^9 +
    1.4753098538474673*10^44*x^10 + 2.0559346241514986*10^44*x^11 +
    1.6882835625514964*10^44*x^12 + 7.606716110341735*10^43*x^13 +
    1.5305410764531565*10^43*x^14 + 5.057848554370418*10^41*x^15)
    /(2.2631962720406472*10^39 + 2.0747988596140944*10^42*x + 5.410284341866251*10^43*
    x^2 + 2.0632316050207164*10^44*x^3 + 3.521034814608054*10^44*x^4 +
    3.2167460431516697*10^44*x^5 + 1.6542542175379038*10^44*x^6 +
    5.511669825549032*10^43*x^7 + 3.371302388006769*10^43*x^8 +
    7.406895515280156*10^43*x^9 + 1.5593809926805384*10^44*x^10 +
    2.1114757191969035*10^44*x^11 + 1.7098739295250467*10^44*x^12 +
    7.645382735089672*10^43*x^13 + 1.531788467484943*10^43*x^14 +
    5.057848554370418*10^41*x^15)

%symbolic GRF  for Figure S7, d):
GRF_d(x) = (6.0961526350200845*10^34*x^4 + 7.74101902589086*10^37*x^5 +
    1.746203723208289*10^40*x^6 + 5.115132287315065*10^40*x^7 +
    3.1895943228064667*10^41*x^8 + 3.342213832117215*10^42*x^9 +
    6.902838350084463*10^43*x^10 + 4.924162063877022*10^44*x^11 +
    3.366474658922477*10^44*x^12 + 6.554770420343171*10^43*x^13 +
    2.4957569986577326*10^42*x^14 + 4.044742151540202 3*10^40*x^15)
    /(1.0922466554269185*10^26 + 3.2780253976670626*10^29*x + 3.43850427345836*10^32*x
    ^2 + 1.5909856119850098*10^35*x^3 + 3.729776750323072 5*10^37*x^4 +
    3.812763936807337*10^39*x^5 + 2.6485070922230037*10^40*x^6 +
    7.472871443792891*10^40*x^7 + 3.848090997460671*10^41*x^8 +
    3.5427300205353945*10^42*x^9 + 6.934872179139399*10^43*x^10 +
    4.925907865407594*10^44*x^11 + 3.3668131964003985*10^44*x^12 +
    6.55495440751056*10^43*x^13 + 2.4957833417073095*10^42*x^14 +
    4.044742151540202 3*10^40*x^15)
```
